# An Analytic Framework for Inferring Population Dynamics from Aggregated Calcium Fluorescence

**DOI:** 10.1101/2024.10.18.619111

**Authors:** Merav Stern

## Abstract

Capturing and inferring brain-wide neural activity remains a significant challenge. Wide-field imaging techniques offer a solution in the shape of simultaneous recording of neural activity across large cortical surfaces at high temporal resolution. Nevertheless, the broad field of view introduced by wide-field imaging limits its spatial resolution, as each camera pixel in a wide-field setup integrates calcium-dependent fluorescence signals from many neurons. Furthermore, calcium indicators that convert neural activity into light emissions distort the neural activity by their dynamics. The inherent noise in recordings, combined with the low spatial resolution and the distorted dynamics introduced by the calcium indicators, makes it particularly challenging to infer underlying neural activity from wide-field fluorescence data. Despite its importance, a rigorously studied analytic solution for this inference problem in the wide-field context has not yet been established.

In this study, we present an analytic solution to the inference problem posed by wide-field imaging. We formulate an optimization problem that establishes a relationship between the biological quantities and the properties of the inferred solution, thereby elucidating biologically interpretable results. The analytic solution we find provides significant advantages, including rapid and accurate inference. Furthermore, we introduce a novel approach to parameter tuning within the optimization framework, which leverages the extensive datasets typical of wide-field imaging. The results demonstrate that our solution surpasses a previously used method for this inference in both accuracy and efficiency. We rigorously validate our solution through comprehensive simulations, large-scale biophysical modeling, and parallel recordings of fluorescence and spiking activity. Collectively, these analyses provide a robust foundation for future applications of our proposed analytical inference.

## 1 Introduction

Optogenetic tools utilize calcium indicators to bind neuron-initiated spikes with fluorescence emission. The resulting fluorescence can then be captured by a camera, making neural activity visible and hence possible to detect.

In the wide-field setting, a vast amount of fluorescence emitted from neural activity is captured by a standard one-photon camera across a large field of view (Cardin et al. 2020). A typical example of such a wide-field view is the entire dorsal surface area of a mouse brain (Silasi et al. 2016). The wide-field setting also enables fast, simultaneous recordings across the whole surface (Gilad 2024). Consequently, wide-field recordings can reliably depict the time evolution of fluorescence signals across all recorded brain areas. However, due to the large field of view, resolution is limited; each pixel captures a light trace that reflects the combined activity of thousands of neurons. In turn, the vast amount of fluorescence captured in each pixel enables recordings to be made through the skull of animals such as mice (Silasi et al. 2016). This, in turn, allows the brain to remain intact during wide-field imaging, making this technique minimally invasive. Because of its low invasiveness, wide-field imaging can be easily used to record awake behaving animals across multiple sessions over several days, enabling the comparison and analysis of recordings across sessions (Musall et al. 2018, MacDowell & Buschman 2020). The long list of advantages of the wide-field method contributes to its popularity as a recording technique, despite its limited spatial resolution.

Deconvolving recorded fluorescence traces in a wide-field setting into their underlying total neural activity is a significant mathematical challenge. The limited spatial resolution, a distinct characteristic of the wide-field method, combined with the presence of noise, necessitates a specialized solution to extract such underlying total neural activity. Many methods exist for deconvolving single-neuron-generated fluorescence recordings into the underlying single neurons spiking activity (Jewell et al. 2019, Friedrich et al. 2017, Pachitariu et al. 2018, Pnevmatikakis et al. 2016, Berens et al. 2018, for example). However, as we demonstrate in this work, the mathematical challenge that stems from the need to infer the total spiking activity of many neurons is unique and solutions designed for the single-neuron scenario do not hold, requiring the development of a new tailored deconvolution method.

Deconvolution is a challenging problem across many disciplines beyond neuroscience, each of which imposes unique requirements that influence the mathematical formulation, constraints, and methodological approaches specific to the field. Over the years, numerous mathematical methods have been developed (Chaudhuri et al. 2014), either to refine existing techniques or to address emerging problems (Engl & Groetsch 1987, Vogel 2002). A prominent example is image processing (Wipf & Zhang 2014), where issues such as blurs, distortions, reflections, or missing data require corrections achieved through deconvolution, typically implemented using iterative algorithms (Bertero et al. 2021).

The Lucy-Richardson deconvolution is an iterative algorithm originally developed to deblur images acquired through a series of lenses that filter the images and introduce noise. In this study, we employ an adaptation of the Lucy-Richardson algorithm for temporal deconvolution of wide-field fluorescence imaging achieved by replacing its filter with temporal calcium dynamics (Wekselblatt et al. 2016). This method serves as a benchmark for comparison with our novel solution introduced in this work, as it represents the only previously published attempt to deconvolve wide-field fluorescence traces.

Directly inferring neural activity from wide-field fluorescence recordings by inverting (convolving) the influence of calcium dynamics is simply impractical. While the relationship between neural activity and calcium indicator dynamics is well known (Zhang et al. 2023, Chen et al. 2013, for example), recovering neural activity from fluorescence requires a derivative operation (since the reverse direction, the calcium levels, integrates a neuron’s spiking activity). This process of direct inverse (derivative-based) operation amplifies noise, making it difficult to distinguish between noise and underlying neural activity. Hence, naive approaches to infer neural activity from fluorescence recordings by directly inverting calcium dynamics are ineffective and require more in-depth mathematical solutions.

To a first approximation, the calcium concentration depicted by fluorescence reflects the volume of recent neural activity. However, neural activity encodes extensive information about brain functions, and hence, it is frequently the detailed information we seek. Therefore, accurately inferring temporal changes in neural activity beyond first-order measurements is significant. Deconvolution methods that leverage calcium indicator dynamics enable inference of relative changes in neural activity over time from wide-field fluorescence data. Inferred fluorescence thus provides a more accurate representation of the temporal evolution of neural activity than raw fluorescence signals alone. Moreover, inference reduces noise based on biologically relevant criteria related to calcium-binding properties, rather than using generic image-processing filters that rely on visibility criteria. It is therefore possible that inference reduces the risk of misclassifying neural activity as noise (or vice versa) and better preserves the integrity of the spiking rate information embedded in the fluorescence data.

Identifying the correct time evolution of neural dynamics via inference can also improve the accuracy of results when conducting more advanced analyses of the data. Such analyses include comparing two brain areas by examining how their neural activities correlate over time (among themselves or with behavioral measures). Other correlation-based techniques, including Principal Component Analysis (PCA) and Non-negative Matrix Factorization (NMF), can also benefit from improved temporal dynamics. These methods are particularly sensitive to the timing of relative changes between variables, whether those variables are neural spiking rates or behavioral measures. This further emphasizes the need to improve the temporal inference of neural activity.

This paper makes multiple contributions toward this goal. First, it presents pioneering work that formulates the inference problem arising from wide-field recordings and provides an analytic solution to it. To the best of our knowledge, only a few prior studies have attempted to infer the activity of multiple spiking neurons from their aggregated fluorescence (Wekselblatt et al. 2016, O’Rawe et al. 2023, Stern et al. 2020). None among these studies presented an analytical solution capable of retrieving a continuous inferred spiking-rate signal. Such analytic solution is crucial for ensuring that inference is fast and efficient while remaining accurate, particularly given the typically large size of wide-field datasets.

We evaluate our solution against the adaptation of the Lucy-Richardson algorithm (Lucy 1974) for wide-field imaging as presented by Wekselblatt et al. (2016) For completeness, we also compare our results to the naive approach of using the fluorescence signal directly as an estimate of neural activity.

An additional contribution we make in this work is leveraging a biophysically detailed volumetric model (Song et al. 2021) to evaluate deconvolution of aggregated fluorescence signals. In this model, neurons exhibit realistic morphologies, calcium concentrations are calculated according to molecular properties, and fluorescence simulations incorporate light propagation through tissue. While this framework has previously been used to investigate single-neuron properties, we adapt it here to generate and evaluate fluorescence signals recorded in a simulated wide-field setup, along with its corresponding underlying neural activity. This dataset enables us to rigorously evaluate the inference performance of deconvolution methods for wide-field imaging in a realistic yet fully known setting.

We also validate our approach on real wide-field recordings, demonstrating its practical applicability to experimental data. For completeness, we verify our method using simulations generated from our model as well.

We make an additional pioneering contribution by introducing a novel parameter-tuning approach. Wide-field recordings provide vast amounts of data, which we exploit for comparing the dynamical properties of the recorded signals with those of the inferred signals obtained using different parameters, thereby identifying optimal parameters. To our knowledge, this constitutes a unique strategy for parameter tuning in inference problems.

This paper hence includes the following content: First, we phrase the mathematical challenge of inferring the total number of spikes, i.e., the population spiking rate, generated by multiple neurons from their collective fluorescence trace. We then present our analytic solution and provide a proof. The performance of our method is benchmarked against a previously applied inference technique for wide-field imaging using synthetic self-generated data, a biophysically detailed dataset, and real recorded data that includes fluorescence (recorded using the wide-field method) alongside spike count recordings. Lastly, we consider a typical experiment in which only fluorescence is recorded and discuss our original parameter tuning in this context.

## 2 Model

We formulate a model to establish the relationship between a recorded fluorescence trace and the collective activity of a large neuronal population from which it arises. To do so, we link the observed fluorescence *y* to the underlying calcium indicator concentration *c*, recognizing that fluorescence provides a noisy measurement of calcium. In turn, we relate the calcium concentration to the neural activity *r*, identifying that neural activity increases calcium concentration, which otherwise decays. These allow us to phrase the relationship between observed fluorescence and aggregated neuronal activity via the following autoregressive model:

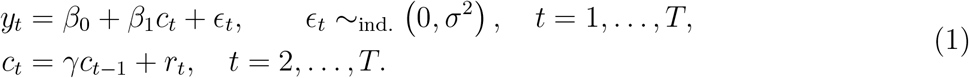

In the model described above, the observed fluorescence at timepoint *t*, denoted as *y*_*t*_, is determined by the average calcium concentration across the neural population at that timepoint, *c*_*t*_ up to some factor *β*_1_, a constant shift, *β*_0_, and noise *ϵ*_*t*_, which is assumed to be Gaussian and i.i.d.. In other words, the fluorescence is a noisy linear readout of the calcium concentration. The calcium concentration at each timepoint, *c*_*t*_, is the result of its decayed value from the previous timepoint, *c*_*t*−1_, by a factor *γ* ∈(0, 1) and the result of the neural activity *r*_*t*_.

In Supplementary Information A, we demonstrate that our model for recorded fluorescence driven by the activity of numerous neurons follows directly from the single-neuron autoregressive model (Vogelstein et al. 2009, Jewell & Witten 2018, Friedrich et al. 2017, Pnevmatikakis et al. 2016). This well-established model relates spiking activity of a single neuron to its resulting fluorescence. Deriving our model from this foundation, clarifies the meaning of *r* in our aggregated model (1). We find that 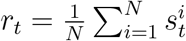, where 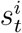 is the spike count for neuron *i* in time bin *t* (where each 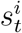 typically takes a value of zero or one). Hence, in our model, (1), *r* is the average number of spikes per neuron in the population of *N* neurons that contribute to the recorded fluorescence during each time bin. We therefore call *r* the *population spiking rate*. This term quantifies the average spiking activity across all neurons that contribute to the observed fluorescence. For brevity, we also refer to *r* simply as the spiking rate throughout the manuscript.

We now choose *β*_1_ = 1 without loss of generality. This corresponds to scaling the calcium by a constant factor, thereby setting a fixed ratio between the spiking rate and the fluorescence. This choice is relevant when analyzing fluorescence and spiking rate recorded simultaneously, as discussed in Section 5.4. In this section, we will explain how to scale the fluorescence in accordance with this choice.

## 3 Optimization

We aim to determine the underlying spiking rate *r* from a fluorescence trace *y* recorded in a widefield setting, utilizing our model (1). We adopt and adapt the approach of formulating and solving an optimization problem to find spikes from a fluorescence trace generated by a single neuron (Jewell & Witten 2018, Friedrich et al. 2017). Since both models, (1) and the single neuron model (see details in Sup Info A), share similar structures, the optimization problem for single neurons can be naturally applied to our model (1) with its corresponding parameters. This leads to the minimization problem that we wish to solve:

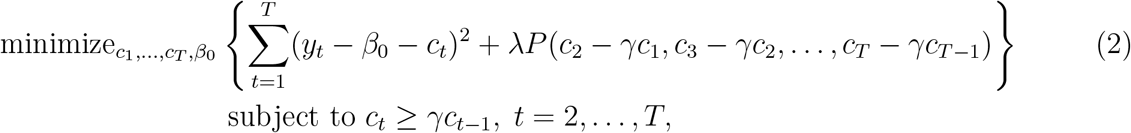

where *λ* is a weight tuning parameter for the penalty *P*.

For convenience, we rephrase (2) in terms of the spiking rate *r*_*t*_. To this end, we define *r*_1_ ≡*c*_1_. We emphasize that this is merely a definition; the spiking rate at *t* = 1 cannot be recovered, and hence *r*_1_ remains biologically meaningless. Using this definition and our model (1), we can express the relation between the calcium concentration and the spiking rate in matrix form as *r* = *Dc*, where *D* is a *T × T* full-rank matrix with 1’s along the diagonal and − *γ*’s just below the diagonal. Since r = Dc is a one-to-one mapping, we can equivalently reformulate optimization problem (2) as follows:

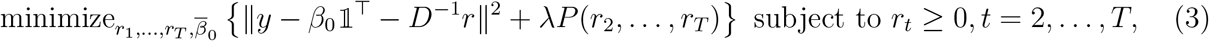

where 1^T^ is a column vector of length *T* with all its elements equal to 1.

At this point, we need to determine the form of the penalty *P* (*r*_2_, …, *r*_*T*_). This choice will dictate the properties of the solution *r* that we derive. Hence, to best fit it for wide-field imaging, we need to consider the characteristics of the underlying population spiking rate *r* in this context. The penalty that was chosen for a fluorescence trace resulting from the activity of a single neuron is no longer relevant. This is because the properties of the number of spikes of a single neuron differ significantly from the properties of the average number of spikes of a large population. The typical profile of the number of spikes per bin in a single neuron consists of many zeros and otherwise features low positive integers. To obtain solutions with such characteristics, *l*_0_ penalty has been chosen (Jewell & Witten 2018, Jewell et al. 2019) or *l*_1_ (Vogelstein et al. 2009, Pnevmatikakis et al. 2016).

In contrast, the spiking rate *r* in our model (1) is a positive continuous variable. This is because it represents the average number of spikes across the entire neuronal population contributing to the fluorescence, and this population can be immensely large; a single fluorescence trace in wide-field recordings aggregates signals from thousands to hundreds of thousands of neurons. As a result, we can treat the population spiking rate *r*, which is the average number of spikes per neuron, as a continuous variable for any practical purpose. Moreover, as an average over such a vast population, we expect it to vary gradually over time while remaining within the physical limits of possible firing rates (see Figure 5(a)i for example). To ensure that the inferred spiking rate *r* exhibits these characteristics, we impose an *l*_2_ penalty on its temporal differences: *P* ((*r*_3_ − *r*_2_)^2^, (*r*_4_ − *r*_3_)^2^, …, (*r*_*T*_ − *r*_*T* −1_)^2^). This penalty enforces the solutions of *r* to be smooth and continuous. An additional benefit of this penalty is that it is differentiable, thereby enabling us to derive and analyze an analytic solution for the spiking rate. Thus, introducing the spiking rate as a population-level measure marks a significant departure from single-neuron models, affecting the optimization problem, the solution methods, and the resulting aggregate neural dynamics solutions themselves.

We hence obtain the following optimization problem for inferring spiking rates from wide-field fluorescence recordings:

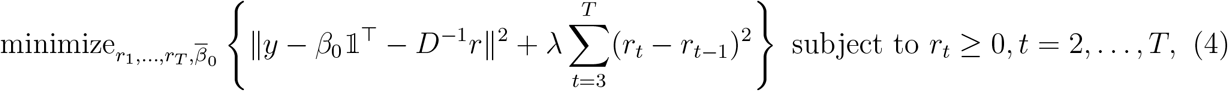

## 4 Solution

At first glance, the optimization problem (4) appears to be a direct sum of two *l*_2_ terms, which suggests it can be solved naively by setting its derivatives with respect to *β*_0_ and *r* to zero and finding their minimum (similar to the Least Mean Squares method). However, a more careful examination reveals that the solution for *r* must be non-negative. Moreover, with *T* fluorescence measurements, only *T* − 1 time points for spiking rates can be determined, since they depend on changes in fluorescence through calcium dynamics (remember that *r*_1_ is merely a notation and does not represent a spiking rate). In other words, we have one degree of freedom, which our solution must “absorb” while preserving its structure to yield a biologically meaningful spiking rate solution. In this section, we prove that this degree of freedom corresponds to an arbitrary constant shift in the fluorescence (a consequence of the Δ*F/F* procedure), which results in the spiking-rate solution also being invariant under such constant shift. We use this degree of freedom to ensure that the spiking rate solution is non-negative, as required.

We begin by simplifying the optimization problem (4). We take its derivative with respect to *β*_0_ and set it to zero, yielding a solution for *β*_0_ as a function of the spiking rate:

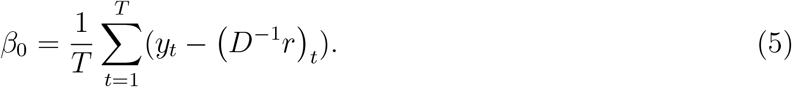

We substitute this expression for *β*_0_ into (4), resulting in the following form of the optimization problem:

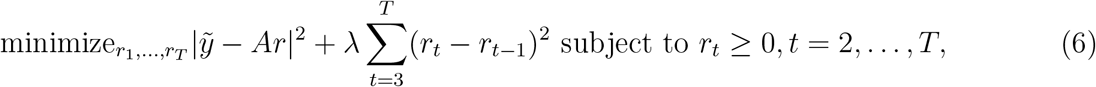

where 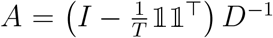 and 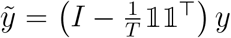, with *I* the *T × T* identity matrix and ]_]_^T^ a *T × T* matrix of ones.

Note that 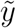 is the mean subtracted version of *y*, and *Ar* is the column-wise mean subtracted version of *D*^−1^*r*. This corresponds to the constant shift of the recorded fluorescence *β*_0_ being arbitrary with respect to the spiking rate. In what follows, we explicitly prove that it makes the spiking rate invariant under a constant shift.

In Supplementary Information B, we provide a proof that the solution to (6) is equivalent to the solution to (4), i.e. the pair 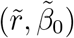 solves (4) if and only if 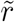 solves (6) and 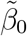 is given by (5). We can now solve the optimization problem (6) by solving (6) to obtain the spiking rate, and then obtain *β*_0_ from its closed-form expression (5). Finding *β*_0_ enables us to reconstruct the noise-removed version of the original recorded fluorescence, along with its shift, which is helpful for testing the results.

To solve (6), it is constructive to present the following, even simpler optimization problem that retains the same *l*_2_ expressions but does not include any constraints on *r*:

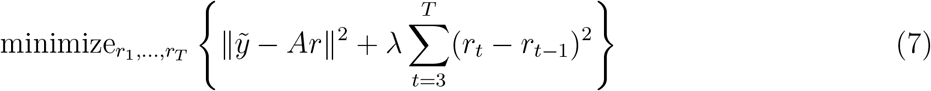

Th is problem can be solved analytically by computing ∇_*r*_*f* (*r*) = 0, with 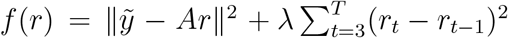 . For convenience purposes, we define *Z* as follows:

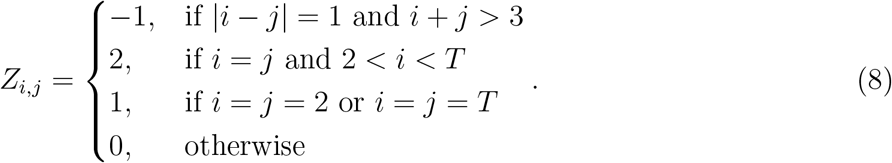

Note that 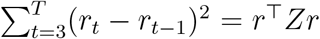 .Therefore, *f* takes the form 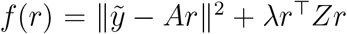 and ∇_*r*_*f* (*r*) = 0 yields the following:

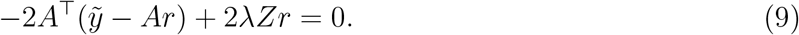

The solution to (9), and hence to problem (7), takes the form

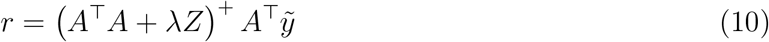

where *A*^T^*A* + *λZ* ^+^ is the pseudo-inverse of *A*^T^*A* + *λZ*. Note that we have absorbed into this pseudo-inverse the degree of freedom that led us to use the mean-subtracted version of the fluorescence, 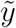, and the column-wise mean-subtracted version of *D*^−1^*r, Ar*. Next, we find the explicit form of this degree of freedom.

We consider 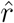, a solution to (7). With it, we can construct a new spiking rate 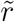 shifted by a constant number *d* by defining 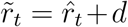 for any *t* = 2 … *T* ; remember that *r*_1_ is merely a notation and does not represent a spiking rate. This construction preserves all the temporal differences of 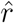, i.e., 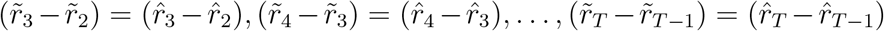 In Supply Information C we prove that 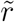 also solves the optimization problem (7) with 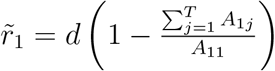

The above implies that problem (7), and consequently the original problem (4), has multiple solutions and is not strictly convex. The solutions differ from each other by a constant shift in the spiking rate *r*_2_, …, *r*_*T*_ and a corresponding adjustment, compensating for that shift, in *r*_1_; recall that *r*_1_ has no biological meaning. As a result, the solution for *β*_0_, equation (5), changes accordingly to accurately relate the shifted spiking rate (through the calcium concentration) to the recorded fluorescence. This concludes our proof that the degree of freedom in the problem translates to the spiking rate solution being invariant under constant shifts.

Solution (10) to problem (7), together with the preceding discussion (formal proofs to it can be found in Supplementary Information B and C), leads to Algorithm 1 for solving (4).

### Algorithm 1

Continuously-Varying Spiking Rate Inference: Solving (4)

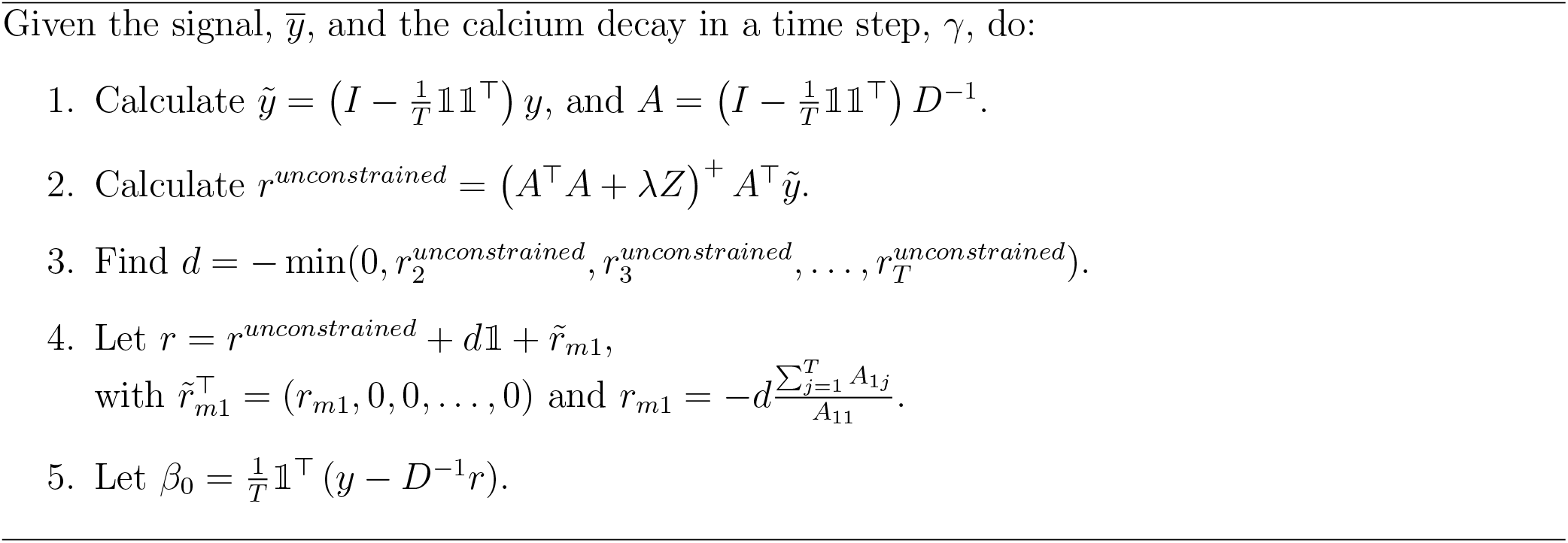

## 5 Results

### 5.1 Evaluation

Here, we evaluate the performance of our Continuously Varying Inference Method (Convar for short), as described in Algorithm 1. To this end, we simulated spiking rate traces (see example in Figure 2(a)i, ivory line; see Supporting Information D for details) and emulated the wide-field recording of fluorescence traces resulting from these spiking rates following our model (1). This process included transforming the simulated spiking rate traces into calcium traces according to *c* = *rD*^−1^ (a direct result of (1), see example in Figure 2(a)ii) and adding Gaussian-independent noise and a constant shift to the calcium traces to generate the recorded fluorescence (Figure 2(a)iii). For the calcium indicator dynamics, we selected *γ* = 0.95, which closely matches its observed values (see Section 5.4).

**Figure 1:**
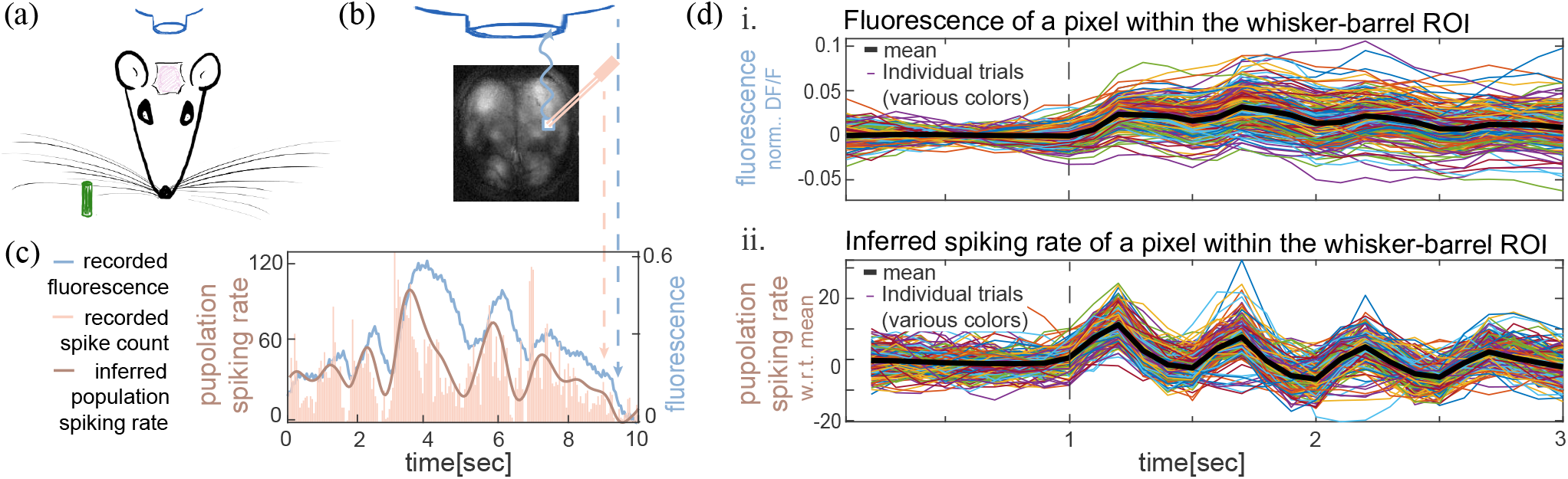
Population spiking-rate inference from wide-field fluorescence recordings. (a) Schematic of the wide-field imaging setup, including whisker stimulation. (b) Example of a wide imaging field and placement of an electrophysiological probe for simultaneous fluorescence and spike count recordings; configuration as in Clancy et al. (2019) see Section 5.4 for details. (c) Example recording obtained using the setup shown in (b). Recorded fluorescence (blue), simultaneously recorded spike counts (ivory bars), and the population spiking rate inferred from the fluorescence using our Continuously-Varying algorithm (brown). (d) i. Normalized fluorescence from a pixel within the whisker-barrel region across 200 trials (colored traces), together with the trial-averaged response (black). Whisker stimulation was delivered at *t* = 1sec (black dashed line). The mean fluorescence contains both sustained and oscillatory components and peaks at approximately 1.75sec. ii. Spiking rates inferred from the same trials, with the mean inferred rate shown in black. The inferred response is predominantly oscillatory and reaches its first peak at approximately 1.25sec. Unpublished data from the Zagha laboratory, UC Riverside; GCaMP6s recordings acquired at 10 Hz.

**Figure 2:**
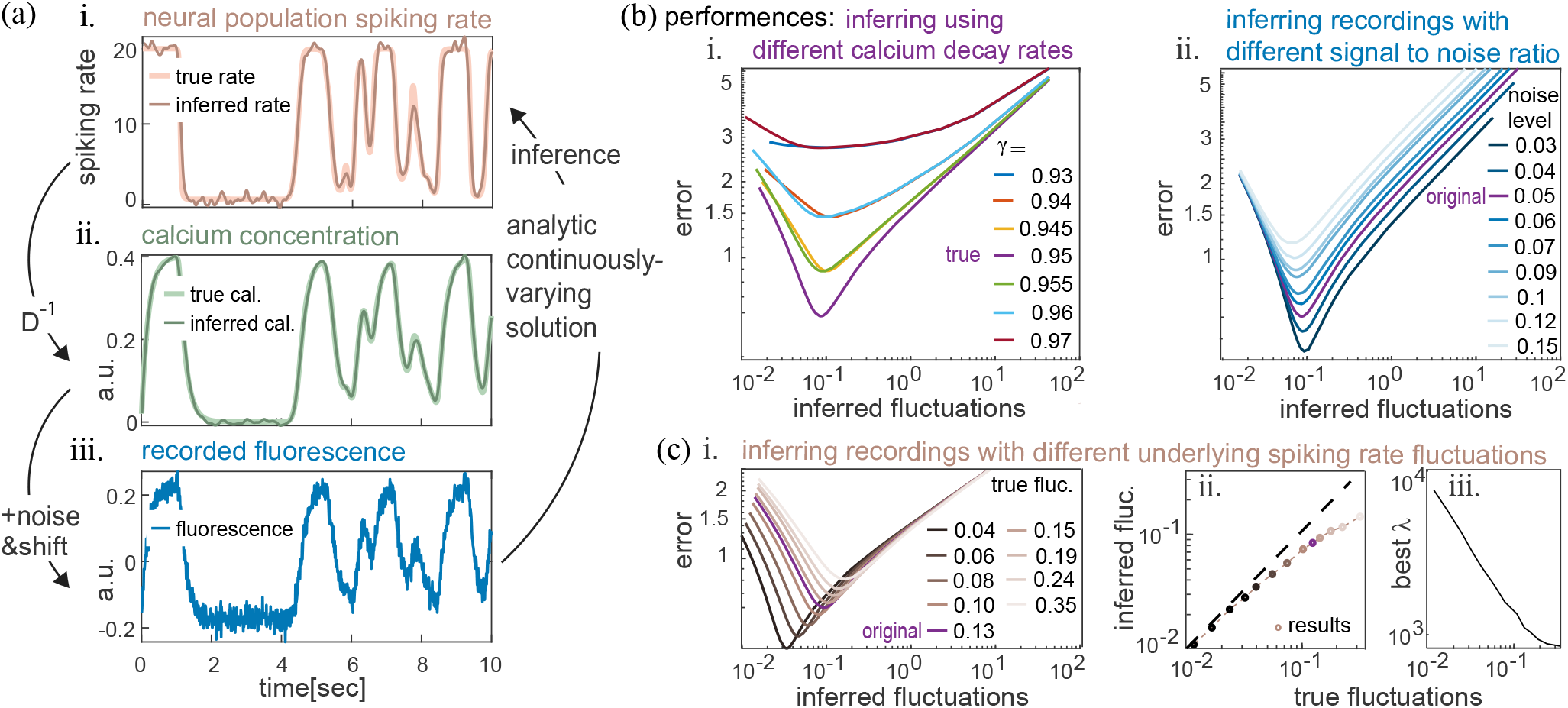
Evaluating the Continuously-Varying Inference, Algorithm 1 (Convar). (a) An example: A simulated underlying spiking rate (i. ivory line) is convolved into calcium concentration (ii. light green line) and transformed into recorded fluorescence (iii. blue line) using our model (1). The fluorescence is then inferred back into spiking rate using Convar (i. brown line). Note that the fluorescence units are scaled by 1*/*1000 to match typical camera settings. (b) i. The mean error (11) between simulated and inferred spiking rates as a function of the mean inferred spiking rate fluctuations (12). The inferred spiking rates were obtained by applying Convar to fluorescence traces generated from simulated spiking rates (as in (a)). For inference, Convar was applied multiple times with different values of *γ* (0.93≤ *γ* ≤ 0.97, yielding multiple lines) and *λ* (10000 ≤ *λ* ≤ 1, yielding variations in the inferred spiking rate fluctuations). Means were taken over 10000 traces, each 10sec long, generated as described in Supporting Information D. Calcium traces were generated with *γ* = 0.95, and for the fluorescence we applied 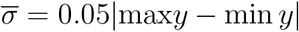. ii. The mean error (11) between simulated and inferred spiking rates as a function of the mean inferred rate fluctuations (12) for different datasets with different noise levels 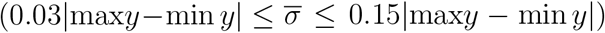. *γ* was held fixed at 0.95 for inference; all other parameters are as in i. (c) i. The mean error (11) between simulated and inferred spiking rates as a function of the mean inferred spiking rate fluctuations (12) for multiple simulated spiking rate sets with different mean (true) fluctuations (see Supporting Information D). *γ* was held fixed at 0.95 for inference; all other parameters are as in (b)i. ii. Mean inferred spiking rate fluctuations as a function of the mean simulated (true) spiking rate fluctuations at the minimum error point identified for each true spiking rate magnitude of fluctuations in i. iii. Values of best penalty *λ* which yield the minimum error points in i., as a function of the true spiking rate fluctuations. The variation in the optimal penalty value allows for fitting the inferred spiking rate fluctuations to the true spiking rate fluctuations (shown in ii.) while maintaining a similar minimal error (in i).

We used Convar to infer the spiking rates from the simulated fluorescence recordings. To assess the quality of the inference, we calculated the following error:

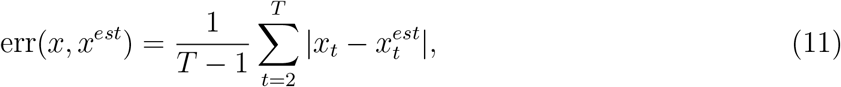

defined as the difference between the original simulated underlying spiking rate, *x* = *r*, and the inferred spiking rate *x*^*est*^ = *r*^*est*^.

We used the mean subtracted versions of both *r*_*t*_ and 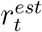, for *t* = 2, …, *T*, to assess the error (11). We chose this approach because the inferred spiking rate is invariant under a constant shift, as implied by the discussion in Section 4.

The magnitude of fluctuations in a trace offers another useful measure for evaluating the quality of our inference. We quantify these fluctuations as follows:

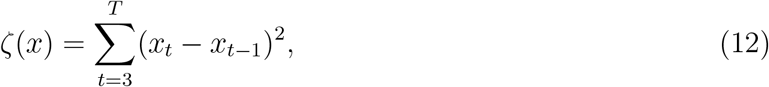

where *x* can be any time-dependent parameter (spiking rate *r*, calcium *c* or fluorescence *y*). The magnitude of fluctuations measures the extent to which the signal changes over time. Note that for *x* = *r*, it is equal to the penalty term in the optimization problem (4). Since the parameter *λ* controls the weight of this penalty, lowering *λ* weakens the penalty, allowing larger fluctuations in the inferred spiking rate, while increasing *λ* strengthens the penalty and suppresses fluctuations. Figure 2(b)i was generated by varying the penalty weight *λ* and the calcium decay constant, *γ*, used for inference. The results were plotted for each value of *γ* according to the fluctuations observed with each *λ* (on the *x*-axis, with higher *λ* promoting smaller fluctuations and vice versa).

For each calcium decay constant *γ* used for inference, there is an optimal level of fluctuations corresponding to an optimal *λ* that produces a solution with minimal error. Convar preserves this optimal level of fluctuations close to its value at the global minimum (lowest error) across a wide range of *γ* (see Figure 2(b)i; the *x* values of the minimum error points are similar across the different *γ* curves). This finding suggests that slight variations in the calcium decay constant *γ* are unlikely to have a significant impact on the optimal inference solution. Small jitters in *γ* can occur in recordings due to several factors, including substantial changes in firing rates, intrinsic properties of neurons, and high tissue density between neurons, among others. Our results indicate that, despite these jitters, the optimal inferred solution is maintained with a high degree of accuracy.

Importantly, Convar is consistent in the sense that it retrieves global minimum error using the same decay rate *γ* as was used for generating the calcium traces (Figure 2(b)i, purple line). Meaning, it accurately identifies the true underlying calcium decay dynamics. At this optimal, and true underlying, decay rate *γ*, the error is also lower at any level of fluctuations. This is a realization of the algorithm being convex, also with respect to its tuning parameters.

The signal-to-noise ratio (SNR) can vary significantly between experiments. Therefore, it is essential to test the properties of the solution amid various noise levels. We changed SNR by changing the variance 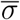 of the noise 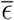 with respect to the total range of the fluorescence. As SNR decreased due to increased noise variance, we found that the minimum error of the optimal inferred solution increased. However, its magnitude of fluctuations remained similar across noise levels (see Figure 2(b)ii; SNR values range from 1*/*0.3 to 1*/*0.15; over this range the minimum error doubles, and the fluctuations increase by about 1.5-fold, despite a five-fold increase in noise variance).

The above suggests that inferred solutions retain their properties amid possible noise variability. This robustness arises because Convar estimates the noise at each time point in the fluorescence recordings and removes it to derive the inferred spiking rate. Hence, noise reduction is a consequential outcome of the inference. Specifically, Convar represents fluorescence as a sum of a calcium signal and noise. It fits the calcium component according to a biologically meaningful criterion, the calcium dynamics, while the remainder of the fluorescence signal is identified as noise and removed. This procedure allows for the extraction of the calcium signal and, in turn, the inferred spiking rate. As a result, Convar solutions are spiking rates with reduced measurement noise and tend to be robust because they are derived with respect to the calcium dynamics.

Inferred solutions by Convar exhibit fluctuations that closely resemble the underlying spiking rate fluctuations (Figure 2(c)ii). This similarity is conserved across an extensive range of fluctuation levels. Convar achieves this with an adjusted optimal penalty *λ*, which leads to the optimal solution (Figure 2(c)i,iii). In other words, the inferred spiking rate fluctuations correctly reflect the underlying spiking rate fluctuations. This enables a proper fit of the inference across various recording scenarios, accommodating a broad spectrum of underlying spiking rate fluctuations.

### 5.2 Comparison

Here, we compare the performances of the Continuously-Varying Inference (Convar for short), as detailed in Algorithm 1, with those of the Lucy-Richardson Algorithm (Lucy for short). To provide a baseline for this comparison, we also assess how Convar and Lucy perform against simply using the fluorescence signal itself as an inferred spiking rate.

Originally, the Lucy algorithm was developed for astrophysics to infer underlying light sources that have been distorted by lenses (Lucy 1974). Recently, it has been adapted to infer spiking rates from fluorescence traces that have been distorted by calcium dynamics in wide-field imaging experiments (Wekselblatt et al. 2016). At their core, the adaptations rely on treating the calcium dynamics as the relevant filter, a temporal filter in this case rather than a spatial filter in the original scenario. This filter can be expressed as *γ*^*t*^ for *t* = 1, …, *p* where *p* is sufficiently large to capture the calcium signal decay. To the best of our knowledge, this is the only algorithm in the published literature that has been used and tested specifically for inferring continuous spiking rate signals from wide-field imaging.

The main advantage of utilizing Lucy for wide-field imaging is its easy, fast implementation. In contrast to our Convar solution, Lucy does not necessarily require parameter tuning. While its number of iterations can be adjusted, our results indicate that the default of 10 iterations consistently performs best across all datasets tested here. Lucy also benefits from pre-written and optimized code in most programming languages. However, since our Convar solution is analytic and not iterative like Lucy, its overall computational efficiency is competitive with that of Lucy.

The main drawback of Lucy is that, at its core, it relies on assumptions that do not necessarily match the characteristics of fluorescence dynamics generated by neural population activity. For instance, Lucy assumes a Poisson noise distribution, which probably does not fit the noise properties in fluorescence recordings. Our tests, which compare the error between the inferred and the underlying spiking rate, reveal that these discrepancies lead to various issues. Most notably, when inferring with Lucy, the correlation between the fluctuations of the inferred spiking rates and those of the underlying spiking rates is lost, as we describe below.

To test Lucy and compare it to Convar, we inferred the same fluorescence traces that we generated in Section 5.1, this time using Lucy.

It is evident that the inferred spiking rate by Lucy does not capture the full span of the underlying spiking rate (Figure 3(a)i, brown vs. ivory lines). This limitation is not critical, as both fluorescence and inferred rates have arbitrary units and thus the range of their values carries no direct biological meaning. Another possible weakness of Lucy is its inconsistency in estimating the calcium decay rate. It achieves minimal error with a decay constant *γ* close to, but not equal to, the true value of *γ*, leading to an incorrect inference of *γ* (Figure 3(b), true *γ* is marked by a purple dashed line).

**Figure 3:**
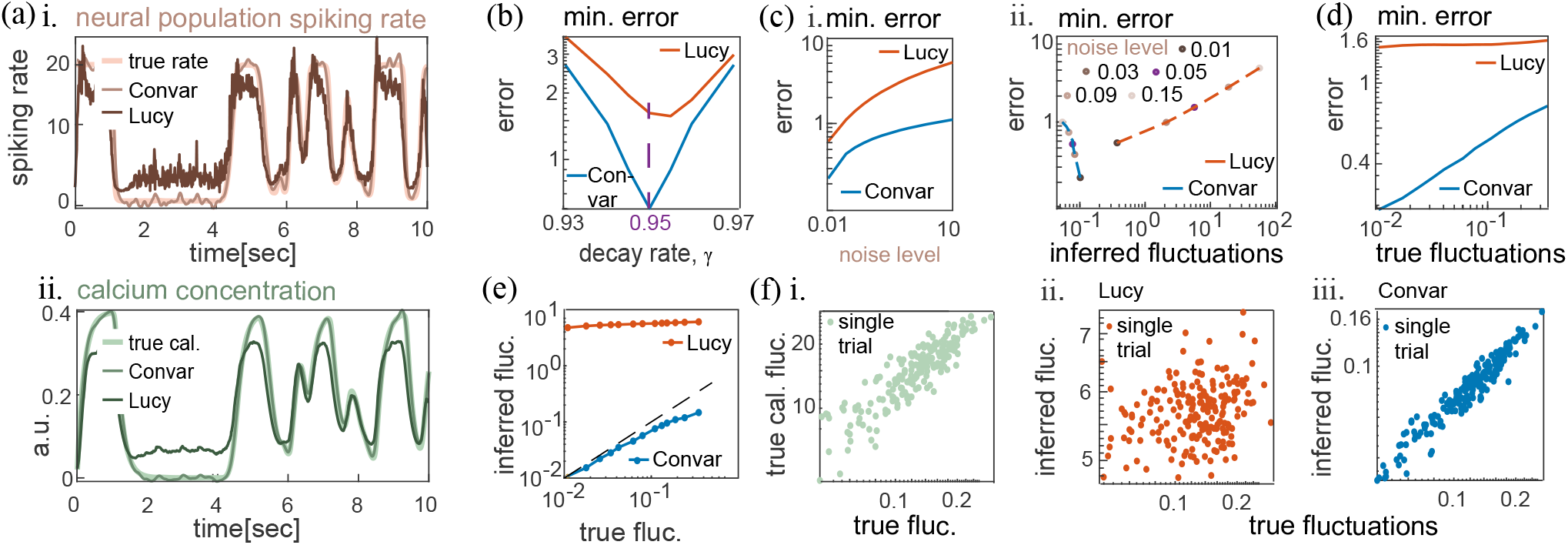
Comparing the Lucy-Richardson Algorithm (Lucy) to the Continuously-Varying Inference Algorithm 1 (Convar). (a) The same example as in Figure 2(a), including additional inference by Lucy. (b) i. The mean error (11) between simulated (true underlying) spiking rates and inferred spiking rates as a function of the different *γ*’s used for inference by Convar and Lucy. For comparison, we also included the error when fluorescence (fluor) is naively considered as the inferred spiking rate. The dataset used in Figure 2(b)i was also utilized here. Similarly, the datasets in Figure 2(b)ii and Figure 2(c) were employed for (d) and (e), respectively. Convar results are represented by the minimum error achieved with each *γ* (minimum error of each line in Figure 2(b)i). The same applies to (d) and (e)i, where Convar minimum errors were derived from Figure 2(b)ii and Figure 2(c)i, respectively. (c) The mean error between simulated (true underlying) and inferred spiking rates as a function of: i. The noise level . ii. The mean inferred rate fluctuations (12) for the noise levels in i. (d) The mean error between simulated (true underlying) and inferred spiking rates for multiple underlying spiking rate datasets with different mean fluctuations. (e) The mean inferred spiking rate fluctuations as a function of the mean underlying spiking rate fluctuations. The black dashed line marks *x* = *y*. (f) For 200 example traces, the relation between the following is presented: the true underlying spiking-rate fluctuation levels and i. the true calcium fluctuation levels, ii.-iii. the fluctuation levels of the spiking rate inferred by ii. Lucy, iii. Convar. The corresponding Pearson correlations (for the displayed examples) are: i. 0.88, ii. 0.55, iii. 0.91. Examples were generated using the best *γ* and *λ* (when applicable) for each inference method.

The spiking rate inferred by Lucy is evidently also noisier compared to the underlying spiking rate (see Figure 3(a)i brown vs. ivory lines). Inferred spiking rates by Convar take a balanced estimate, capturing the changes of the underlying rate more accurately (see Figure 3(a)i thin brown vs. thick brown lines). This discrepancy by Lucy creates local dissimilarities between the underlying and the inferred spiking rates. At many timepoints, Lucy introduces changes in the inferred spiking rate when the underlying spiking rate is constant. As a result, the fluctuations inferred by Lucy remain consistently higher than those of the underlying rate (Figure 3(e)). Lucy’s consistently high fluctuations in its inference solutions suggest that it may be suitable for inferring fluorescence recordings generated by higher fluctuating underlying spiking rates.

The difficulty Lucy has in fitting the fluctuations manifests both on average (Figure 3(e)) and in individual traces (Figure 3(f)ii). Our individual simulated calcium traces exhibit variable fluctuations that are highly correlated with those of the single spiking-rate traces from which they were generated (Figure 3(f)i, Pearson correlation 0.8846). In other words, the calcium data (and the fluorescence as a result) maintain the fluctuation structure of the underlying spiking rate. Lucy infers spiking rate fluctuations that are less correlated with the underlying spiking rate fluctuations (Figure 3(f)ii, Pearson correlation of 0.547). This is not inherent to inference; Convar successfully maintains the structure of fluctuations in the data even at the single trace level (Figure 3(f)iii, Pearson correlation 0.8936). This difference highlights a limitation of Lucy, which is an outcome of its assumptions being less fitting for a wide-field setting. We hence find the fluctuations’ correlation to be an informative metric for evaluating inference quality. We adopt its use, particularly for real data where the underlying spiking rate and thus the direct inference error are harder to estimate.

### 5.3 Anatomical and Optical (NAOMi) Simulated data

In this section, we evaluate inference performed by the Continuously-Varying method (Convar for short; Algorithm 1) and the Lucy–Richardson algorithm (Lucy for short) on fluorescence resulting from the activity of many neurons, generated using the Neural Anatomy and Optical Microscopy (NAOMi) simulator (Song et al. 2021). The advantage of using NAOMi is that it supplies anatomically and biophysically detailed cortical datasets that are independent of our model assumptions (1), while still allowing full access to the ground truth by virtue of simulation.

The NAOMi simulator (Song et al. 2021) constructs a detailed volumetric model of cortex that includes vasculature and neurons, along with their somata, dendrites, and axons. It simulates neural spiking activity and its impact on the calcium concentration by accounting for calcium flux via voltage-dependent channel kinetics. It also simulates light emission and propagation by considering the binding of calcium to fluorescence-emitting molecules and the subsequent propagation of fluorescence through tissue (scattering, absorption, and reflection), which results in realistic optical readouts of neuronal activity.

We simulated a cortical-like volume using NAOMi, measuring 250 *×* 250 *×* 100 (*µm*)^3^. This simulated volume contained 3,959 neurons, each spiking at an average rate of 10Hz. We “recorded” the simulated fluorescence resulting from this neural activity for the equivalent of half an hour and split the results into 60 traces of half a minute each. To emulate single-pixel imaging within a wide-field setup, we calculated the total fluorescence emitted from the entire volume, which is the sum of all photons that would be captured by NAOMi imaging simulator. ^1^

Because neurons differ in their luminescence strength (sometimes by orders of magnitude), as reflected in NAOMi simulations, we needed a scheme to estimate the relative contribution of each neuron to the wide-field fluorescence trace. Hence, for each neuron *i* we defined a weight *α*^*i*^ equal to its baseline fluorescence, reflecting its approximate relative contribution. We used these weights to calculate the weighted spiking rate (see also the discussion in Supplementary Information A), 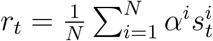, where 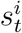 is the spike train of neuron *i*. This weighting allowed Convar and Lucy to achieve successful inference (see Figure 4).

**Figure 4:**
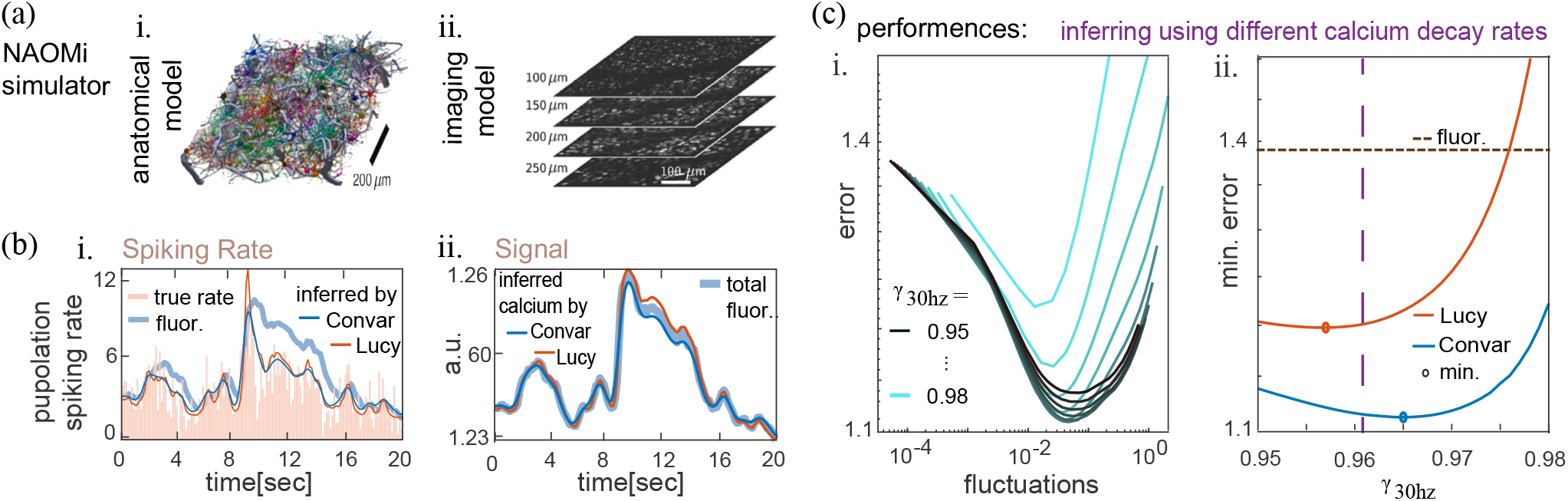
Evaluating inference with biophysical NAOMi simulations. (a) Adapted from NAOMi (Song et al., 2021) i. An illustration of a simulated cortical volume containing vasculature and neurons with dendrites and axons. ii. An illustration of simulated imaging of the activity from the volume in i. To simulate wide-field single-pixel fluorescence, we calculated the total fluorescence captured by the simulated imaging of the entire volume. (b) An example: i. A total population spiking rate calculated from a NAOMi simulation (ivory bars) and the inferred spiking rate by Convar (blue line) and Lucy (red line). ii. The resulting total fluorescence (green line) from the same spiking activity used in i., as simulated by the NAOMi imaging model. The inferred calcium concentration by Convar (blue line) and Lucy (red line). (c) i. The mean error (11) between NAOMi simulated population spiking rates and Convar inferred spiking rates as a function of the mean inferred spiking rate fluctuations (12). For inference, Convar was applied multiple times with different values of *γ* (0.95 ≤ *γ* ≤ 0.98, yielding multiple lines) and *λ* (1000 ≤ *λ* ≤ 0.01, yielding variations in the inferred spiking rate fluctuations). ii. The mean error (11) between NAOMi simulated population spiking rates and the inferred spiking rates by Convar (blue line) and Lucy (red line), as a function of the different *γ*s used for inference. Circles mark minimum errors. For comparison, we also included the error when fluorescence (fluor) is naively considered as the inferred spiking rate (dashed black line). Convar results are represented by the minimum error achieved with each *γ* (minimum error of each line in i.).

Convar consistently produced the lowest error, remaining below Lucy’s minimum error with most decay constants, *γ*, and variable penalty weights, *λ*, tested (Fig. 4(c)i,ii). Both Convar and Lucy captured the fluctuations present in the simulations (Fig. 4(b)), with Lucy producing more fluctuating solutions, consistent with earlier results (Section 5.2, Figure 3(a)i and Figure 3(e)).

Unfortunately, NAOMi simulations generate underlying spiking rate fluctuations that are relatively consistent at the population level, with only about a 5% variance across trials, which hinders our ability to use NAOMi for evaluating how effectively inference methods track fluctuations across different scales. This consistency also leads to weak correlations, approximately 0.2, between the underlying weighted spiking rate and calcium or inferred spiking rates, reflecting noise rather than the ability or inability of the inference methods to follow fluctuations across scales.

Since NAOMi directly simulates calcium flux, binding, and fluorescence emission, it lacks an underlying decay rate *γ*. Song et al. (2021) estimated the effective decay constant for NAOMi, in the setting we employed, which simulates GCaMP6f kinetics at a sampling rate of 30Hz, as *γ* = 0.961 (Fig. 3(c)ii purple dashed line). This aligns with the decay rate reported for GCaMP6f by Chen et al. (2013), who also indicated approximately 10% deviations (Song et al. (2021) did not provide an estimate for these deviations in NAOMi). Our inference achieved minimal error with similar decay constants within this margin, estimating *γ* as approximately 0.957 with Lucy and 0.965 with Convar.

Overall, the results with NAOMi simulations are consistent with those obtained using our model in Section 5.2.

### 5.4 Recorded Data

Next, we tested the performance of the Continuously-Varying Inference (Convar for short, as detailed in Algorithm 1) and the Lucy-Richardson Algorithm (Lucy for short) on real recorded data. We used simultaneous recordings of wide-field fluorescence and spike counts. Unfortunately, it is currently not technically feasible to record the exact underlying neural activity alongside its corresponding fluorescence trace in a wide-field setting. As a result, we do not have the precise underlying spiking rate to test the inference methods. However, we hold an estimate of it. The dataset we use includes spikes recorded from the spatial location of the fluorescence origin from Clancy et al. (2019); see illustration in Figure 1(b)&(c) and Supporting Information E for more details about this dataset. Clancy et al. (2019) verified that the spikes in the dataset decently represent the underlying spiking rates that drive the recorded fluorescence. For our study purposes, this implies that the spiking rate is also a noisy measurement of the exact underlying spiking rate. We will discuss the implications of this as we tune the inference parameters for Convar.

Prior to testing the algorithms on the data, we rescaled the recorded fluorescence traces to ensure that the assumption *β*_1_ = 1 we made in Section 4 is satisfied. Since both the spiking rate *r* and the fluorescence *y* are provided in the dataset, *β*_1_ is implicitly given as well. We calculate its value, denoted 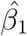, using equation (29) with the same value of *γ* that we use for inference (see Supporting Information F). For inference, we multiply the recorded fluorescence by 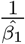 so that the assumption *β*_1_ = 1 in our model (1) is met for the rescaled fluorescence data. This normalization allows a direct comparison between the (mean-subtracted) inferred spiking rate and the (mean-subtracted) recorded spiking rate.

We inferred the data using Convar with various calcium decay constants (*γ* values) and penalty parameters (*λ* values). As we discussed in Section 5.1, any change in the penalty alters the fluctuations in the inferred spiking rate. We plotted the error as a function of fluctuations for each *γ* (see Figure 5(b)i). Our results show that Convar remains convex with respect to the error for both parameters, *γ* and *λ*, consistent with our findings on simulated data (Figure 2(b)i and Figure 4(c)i). For low fluctuations, the error curves are shallow, indicating that small deviations in either *γ* or *λ* have little effect on the inferred spiking rate. This robustness is important because the true underlying calcium decay constant is not precisely known. Clancy et al. (2019) reported *γ* = 0.97 for single neurons in this dataset. Similarly, Chen et al. observed a comparable value with a 10% deviation across cells, *γ* = 0.97 ± 0.003. Such deviations likely reflect natural cell variability, including differences in physical properties and spiking rates (Chen et al. 2013).

In wide-field recordings, deviations can also stem from the presence of non-neuronal tissue, such as glial cells, or from fluorescence that originates from axons or dendrites rather than somas. Convar achieves minimum error when using decay constant values within the range reported for single cells (*γ* = 0.973, Figure 5(b)iii), suggesting that the calcium decay rate observed in single cells is a reasonable estimate for the wide-field setting as well. Lucy yields minimum error with a similar decay (*γ* = 0.974, Figure 5(b)iii).

We further assessed the robustness of our estimate for the calcium decay rate *γ* by identifying the *γ* values that minimize the error for each of the 65 data segments (see Supporting Information E). Our analysis shows that the error between the recorded and inferred spiking rates is minimized at *γ* = 0.9749 ± 0.0042 for Convar and *γ* = 0.9731 ± 0.0047 for Lucy, where ± denotes the standard deviation. These calculated ranges fall within the known single-cell range.

We als o found that Lucy infers higher fluctuations in spiking rates compared to Convar (Figure 5(b)ii), a trend consistent with findings from simulated data (Figure 3(e)). However, the overall range of Lucy-inferred spiking rates is often narrower than that of Convar (Figure 5(a)i), again in agreement with observations from the simulated data (Figure 3(a)i).

To assess the structure of fluctuations, we examined correlations between recorded fluorescence and spiking rate fluctuations. These correlations are inherently limited by noise and the partial representation of the true spiking rate in recordings, resulting in modest observed correlations (Figure 5(d)i, Pearson correlation 0.375). Consistent with simulated data results, Lucy struggles to preserve the observed fluctuation structure (Figure 5(d)ii, Pearson correlation 0.138), whereas Con-Var better retains some of this structure (Figure 5(d)iii, Pearson correlation 0.265).

### 5.5 Analyzing Recorded Fluorescence

All the above analyses directly compared the inferred spiking rate with an underlying spiking rate, either recorded or simulated.

However, in a typical wide-field experiment, only fluorescence data is recorded. Hence, for our algorithm to be practical, we need to be able to use it, including tuning its parameters and testing it, without using underlying spiking data.

Here, we do just that. We tune and test our Continuous-Varying Algorithm by using only fluorescence traces. We then compare our results to the previous section. In other words, in this section, we ignore the recorded spiking rates but eventually utilize the information they carry to evaluate performance.

First, we tune the calcium decay rate *γ*. To do so, we use its known value from single-neuron data. According to our model, equation (1), the calcium decay rate for the calcium concentration of many neurons follows the decay rate of single neurons (see also Supplementary Information A). Our results in Section 5.4 confirm that; they demonstrate that the calcium decay in a wide field setting can be estimated from the single-neuron calcium dynamics.

Our main challenge is to find a penalty weight *λ* to use in the Convar algorithm without relying on minimizing the error between the inferred and underlying spiking rate. We survey here two methods for doing so. The first method was proposed by Jewell & Witten (2018). It divides the fluorescence trace into a training set and a testing set. While this method fits noisy fluorescence recordings, it does not account for noise in the spiking rate.

We present a second, novel approach to fit the penalty weight *λ*. It takes advantage of the vast amount of data available. It maximizes the preservation of data properties (prior and post inference) in relation to other parts of the data. For spiking rate inference, we use the magnitude of fluctuations of the inferred spiking rate (relative to the distribution of the fluorescence fluctuations) as the conserved property. We discuss the two tuning methods for *λ* in the following sections.

#### 5.5.1 Choosing Penalty by Cross Validation

In this section, we summarize the scheme for selecting the penalty weight *λ* as described in Appendix B of Jewell & Witten (2018). To ensure coherence, we include the complete algorithm from Jewell & Witten (2018) in Supporting Information G.

In short, the algorithm divides the fluorescence trace into two subsets, one containing recordings at even-indexed time points and the other at odd-indexed time points (Figure 6(a)i,iii). One subset serves as the training set for inferring spiking rates, while the other functions as the testing set. We assume the fluorescence changes occur slowly relative to the changes in the spiking rate and that noise is independent across time (equation (1)). After inference and convolution back to the calcium concentration, the average of two consecutive points in the inferred calcium concentration is compared with the corresponding point of the fluorescence signal in the other subset prior to inference (Figure 5(a)ii). The penalty weight *λ* is selected as the largest value that yields an error less than the minimum error plus the standard deviation of the error at the minimum error point (Figure 6(b)).

**Figure 5:**
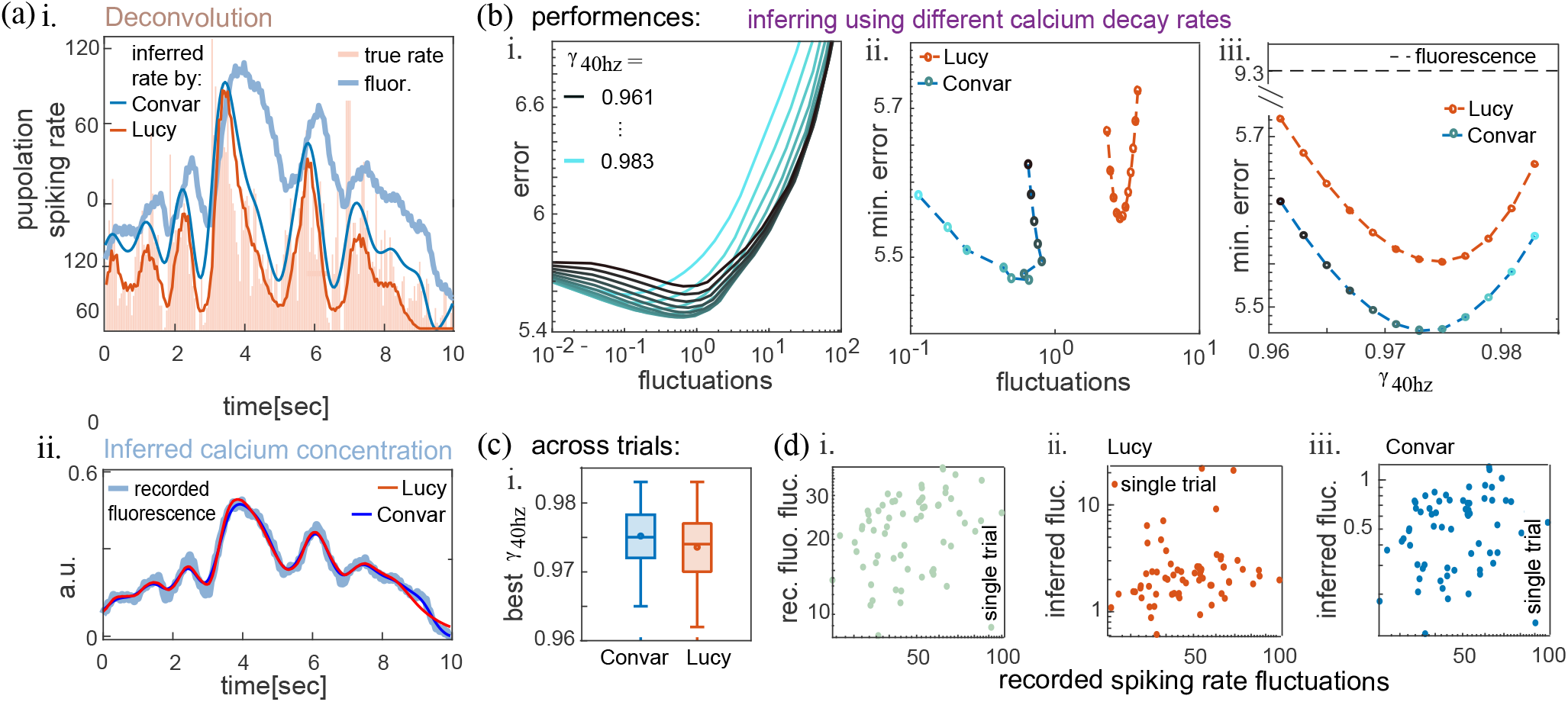
Performance on parallel spike counts and wide-field fluorescence recordings. (a)i. An example of spike counts (ivory) and the inferred spiking rate using Convar (blue line) or Lucy (red line). ii. For inference, the fluorescence recorded in parallel was used (light blue). (b) The mean error (11) between the recorded spiking rates and inferred spiking rates by Convar as a function of the mean inferred rate fluctuations (12). The inferred spiking rates were found by applying Convar on the fluorescence traces recorded in parallel. Convar was applied many times, each time with a different *γ* (yielding the multiple lines) and *λ* (yielding changes in the inferred spiking rate fluctuations), with 0.961 ≤ *γ* ≤ 0.983 and 10000 ≤ *λ* ≤ 0.001. ii. The same as i, including Lucy for inference (red line). Results presented for Convar (blue line) include the minimum error achieved by it with each *γ* (blue circle shades fit the lines in i from which the minimum was taken). iii. Same as ii, but as a function of *γ* used for inference. (c). The medians (lines inside), means (circles), quartiles (boxes), and outliers (whiskers) of all data segments’ best *γ* yielding minimal error between underlying and inferred spiking rate for inference using Convar (blue) or Lucy (red). (d) The recorded spiking rate fluctuations and their corresponding i. recorded fluorescence fluctuations, ii. Lucy inferred spiking rate fluctuations, iii. Convar inferred spiking rate fluctuations. The Pearson correlations are: i. 0.375 ii. 0.138 iii. 0.265. The best *γ* and *λ* according to (b) were used for inference.

**Figure 6:**
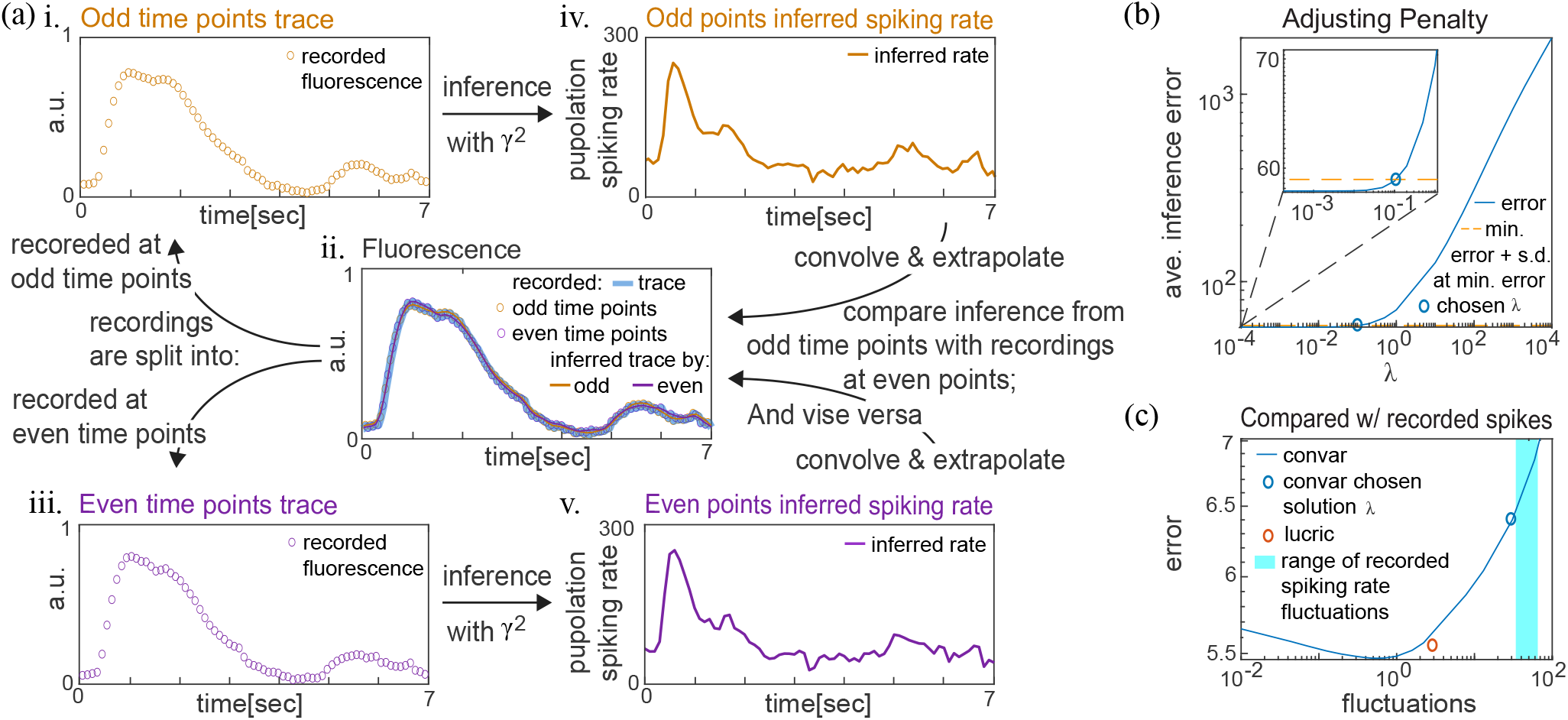
Evaluating the penalty weight *λ* by a cross-validation scheme. (a) An example of a fluorescence trace (ii. light blue), separated into i. odd-, and iii. even-indexed time points. Inferred from them, by Convar, are the iv. odd- and v. even-indexed time points’ spiking rate. Their convolutions into calcium signals are compared with the recorded fluorescence at the even- and odd-indexed time points (respectively; see Supporting Information G for the complete algorithm). This process is repeated with different *λ* values. (b) The average error (11) between the recorded fluorescence at even-or odd-indexed time points and the convolved calcium of the spiking rate inferred from fluorescence at odd- or even-indexed time points (respectively), as a function of *λ*. The chosen penalty (blue circle) is the maximum *λ* with error (blue line) smaller than the minimum error achieved plus the standard deviation at the minimum error (orange dashed line). The mean error (11) between the recorded spiking rates and inferred spiking rates as a function of the mean inferred rate fluctuations (12). The plot is taken from Figure 5(b)i. with *γ* = 0.973. Varying *λ* yields changes in the inferred spiking rate fluctuations. Marked by a blue circle is the *λ* chosen by the cross-validation scheme in (b). The red circle marks the solution given by Lucy for comparison.

We applied this procedure (also described in detail in Algorithm 2, Supporting Information G) to the fluorescence recordings from Clancy et al. (2019). The resulting error between the inferred spiking rate and the recorded spike count, using the penalty weight *λ* determined by this approach, is satisfactory (Figure 6(c), blue circle). Yet, this error does not reach the minimal value achievable with Convar (Figure 6(b), minimum of the blue line). In contrast, the fluctuations of the inferred spiking rate produced by the *λ* selected through this method closely match, on average, the fluctuations in the underlying recorded spike count (Figure 6(c), light blue area). These results indicate that this approach effectively separates noise from the calcium signal, enabling recovery of the spiking rate with dynamical properties that closely correspond to the underlying signal.

#### 5.5.2 Choosing Penalty by Maintaining Signal Properties

We introduce an alternative, novel approach for tuning the penalty weight *λ*. Our approach relies on finding the value of tunable parameters that best preserve some dynamical quantities across the data after inference. By quantifying the relative amount of a dynamic quantity present in each data segment and using that to choose parameters for inference, we leverage the superfluity of information in large datasets. This strategy is well suited to wide-field experiments, where substantial fluorescence data is available. The central idea is to select a penalty weight that preserves the relative amount of a fundamental dynamical quantity after inference.

The rationale for this approach is that the selected dynamical quantity captures a key property of the underlying signal. While inference may alter its absolute value, the relative amounts across segments should remain consistent. In our analysis, we use the magnitude of fluctuations as the primary dynamical quantity. We expect segments with larger fluorescence fluctuations to yield larger inferred spiking rate fluctuations, and conversely, segments with lower fluorescence fluctuations to yield lower inferred spiking rate fluctuations (see Figure 7(a) for an example). Thus, we require the inference to preserve the relationships between levels of fluctuations before and after inference.

**Figure 7:**
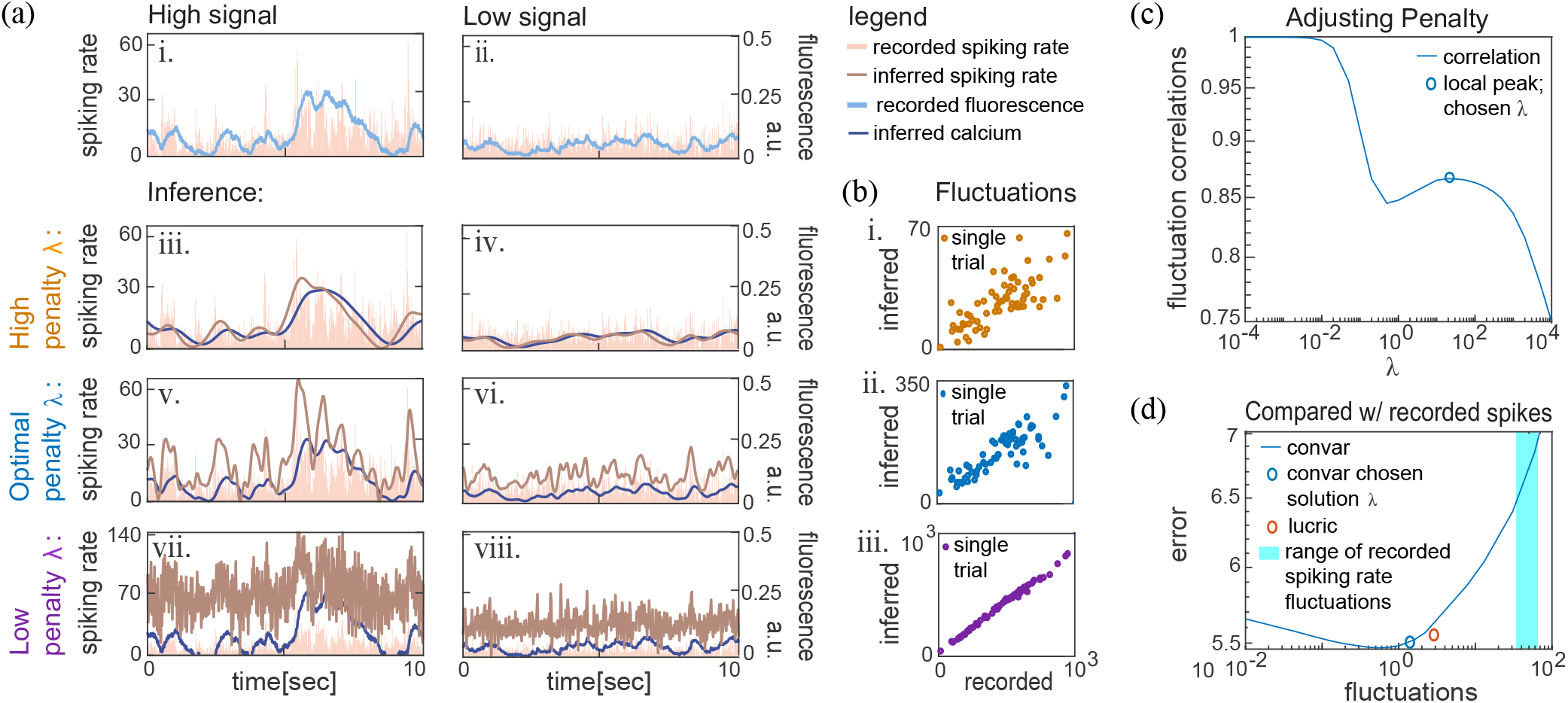
Estimating the penalty weight *λ* according to fluctuation correlations. (a) i,ii. Two examples of recorded fluorescence (light blue) and the parallel recorded spike counts (ivory). iii-vii. The inferred spiking rate (brown) from the two fluorescence examples using iii,iv. high, v,vi. medium and vii,viii. low penalty values and their resulting convolved calcium (blue). The optimal penalty succeeds best at differentiating between the high and low signal examples. (b) Scatter plots of all 65 fluorescence segment fluctuations as recorded and their corresponding calcium segment fluctuations as calculated from the inferred spiking rates convolved back into calcium. Pearson correlation for the plots: i. 0.753 ii. 0.867 iii. 0.997. (c) The correlation between the recorded fluorescence fluctuations and the fluctuations of the convolved calcium from the inferred spiking rate, as a function of *λ*. The chosen penalty (blue circle) locally maximizes the correlation. Same as Figure 6(c) with the blue circle representing the local minimum solution chosen by fluctuation correlations, as marked here in (c).

Maintaining the order and ratio of inferred spiking rate fluctuations relative to fluorescence fluctuations across segments requires selecting an appropriate penalty weight *λ* for the inference. In Convar, the penalty and the fluctuations (12) of the spiking rate share the same mathematical form, as discussed in Section 5.1. Therefore, careful adjustment of the penalty weight *λ* also adjusts the fluctuations, with higher penalty weights leading to lower fluctuations and vice versa. Proper adjustment of *λ* enables accurate estimation of the signal-to-noise ratio in the fluorescence trace, facilitating effective noise removal. High penalty values for *λ* result in low fluctuations, which can cause excessive classification of signal as noise and substantial signal loss (Figure 7(a)iii). Conversely, low penalty values for *λ* lead to high fluctuations, resulting in insufficient noise removal and misclassification of noise as signal (Figure 7(v)iii), producing an overly noisy inference. The goal is to maximize information retention in the spiking rate while optimizing noise removal by preserving the ratio of dynamic quantities, specifically the fluctuations, after inference.

To select *λ*, we calculate the correlation between fluctuations of different segments before and after inference, using the inferred calcium signal, which we obtain by convolving the inferred spiking rate. We perform this calculation across a range of *λ* values. When *λ* is zero, no noise is removed, so the restoration of the signal after inference and convolution is exact, yielding a correlation of 1 between segment fluctuations. However, this setting retains all fluorescence noise in the inferred spiking rate, preventing accurate inference because noise is interpreted as spiking activity. As we increase *λ*, the correlation reaches a local maximum at an intermediate value. At this point, we preserve most of the dynamical information while removing the majority of noise from the signal. We select this value of *λ* as the penalty weight for Convar.

We find that the local-maximum approach described above produces solutions that closely match the best available one (Figure 6(d)). By using fluctuations averaged over segments, rather than point-by-point comparisons as in cross-validation, we account for noise in both the fluorescence recordings and the underlying spiking rates. This method enables us to evaluate the inferred spiking rate while considering both sources of noise. This approach is appropriate for our data, since the recorded spike count in Clancy et al. (2019) is only correlated with the true underlying spiking rate of the recorded fluorescence. In other experimental setups, additional noise in the underlying spiking rate may arise if only a few traces are available to represent an average response or when comparing across multiple experiments.

## 6 Discussion

Many algorithms exist to correct and analyze spatial distortions and features in wide-field recordings. They include methods for correcting small motions in the recorded plane or camera (Reddy & Chatterji 1996, for example), denoising (Buchanan et al. 2018, for example), defining regions of interest (Saxena et al. 2020), and more. While they provide useful tools to reduce the noise and analyze the data, they are not capable of, nor are they intended to, correct distortions in the temporal domain. In other words, after treating spatial distortions, it remains a challenge to portray the neural activity through time correctly. This is because the fluorescence intensity through time is a transformation, in the temporal domain, of the original neural activity magnitude due to the calcium dynamics. Less than a handful of attempts were made to estimate the correct temporal neural dynamics from the fluorescence (O’Rawe et al. 2023, Wekselblatt et al. 2016, Peters et al. 2021, Stern et al. 2020). Here, we offer pioneering work in which we present an analytical solution to the inference problem of recovering continuously varying population spiking rate dynamics from the fluorescence recordings obtained in a wide-field setting.

We compare our new analytical solution of Convar (Algorithm 1), to the Lucy-Richardson algorithm (with its adaptation to the wide-field setting (Wekselblatt et al. 2016); First, we demonstrate that our solution yields significantly lower error than the Lucy-Richardson algorithm in the scenarios we tested, including various SNR levels and underlying fluctuations in synthetic data (Figure 3(b) and Figure 4(c)ii) and real data (Figure 5(b)ii). Second, we demonstrate that our solution is more consistent than the Lucy-Richardson algorithm in the sense that it retrieves the lowest error with the correct underlying calcium decay dynamics (Figure 3(b)). Our Continuously-Varying solution also best maintains fluctuation correlations (Figures 3(f)iii and Figure 5(d)iii).

The solution we bring here also requires parameter tuning. While this increases the initial effort in setting up the inference for each setup, it allows our Continuously-Varying solution to be adaptable, designed for the possibility of being broadly applicable to many setups (including a variety of animals, brain regions, calcium indicators, and more). We study two different approaches for tuning the penalty weight, including a novel scheme for the unique case of noisy spiking rate (with respect to the fluorescence or some other defined criterion). We find that after tuning, even when the underlying spiking rate data is unavailable (as in a typical wide-field experiment), the result is satisfying and competitive with other possible non-tuning-requiring approaches, such as the Lucy-Richardson algorithm. Overall, we find that the Continuously-Varying solution’s lower error and better fluctuation correlation are encouraging for addressing the potential challenges in tuning.

Our optimization problem formalism in equation (4) allows us to develop an inference method that accounts for the shape of calcium dynamics, the bounded amplitudes of spiking rates, and random, time-independent noise as described in equation (1). These considerations enable our approach to minimize inference error while preserving the structure of the data. Our method also yields consistent spiking rate solutions when parameters jitter, a scenario that commonly occurs in experimental settings, as discussed in Section 5.4.

We envision our inference solution becoming part of the “pipeline” for wide-field imaging experiment data analyses. We implement it such that its output, the inferred spiking rate solution, has the same structure as the input, the fluorescence recording. Because the inference operates in the temporal domain and does not alter the data structure, it can be incorporated at any stage of the pipeline. The placement of inference within the pipeline affects noise removal, which is a goal shared by several processing steps. We leave the comparison of the different pipeline results for future studies. We recommend applying our inference solution after spatial distortion corrections and before any dynamics-related analyses, such as dimensional reduction, identification of regions of interest, PCA, NMF, or mixed linear models. With these steps, the inference is expected to perform best, leading to more accurate analysis results altogether.

We make our code available at https://github.com/meravstr/Wide-Field-Inference-2024.

The Supporting Information file includes the derivation of our model from single-neuron models, proofs of the mathematical statements presented in this work, detailed descriptions of the simulated and recorded datasets, an algorithm for finding *β*_1_ for datasets with simultaneous fluorescence and spiking activity recordings, and an algorithm for selecting the penalty parameter *λ* for typical datasets containing only fluorescence measurements.

## Supporting information

sup inf

## Footnotes

1 We are grateful to Alexander Song, the lead author of NAOMi, for his guidance in implementing a computationally efficient wide-field–like recording from NAOMi.

