## Supplementary material for "An Analytic Framework for Inferring Population Dynamics from Aggregated Calcium Fluorescence": sup inf

Merav Stern[1,*]

[1]The Center for Theoretical Studies in Physics and Biology, The Rockefeller University, New York, NY, 10065, USA
[*]

In the Supporting Information file, the derivation of our model from single-neuron models is presented in Section A. Propositions 1 and 2, which prove the mathematical statements presented in this work are presented in Sections B and C, respectively. Section D describes the simulated dataset, while Section E provides details of the recorded dataset. In Section F, we demonstrate how to estimate the coefficient $\beta_1$ from a dataset when it includes both fluorescence and spiking activity. Finally, Section G provides a detailed description of the process for selecting the penalty parameter $\lambda$ using a cross-validation scheme when only fluorescence data is available.

#### A Derivation of Our Population Fluorescence Model from Single-Neuron Models

We begin by examining how the spiking of an individual neuron contributes to a recorded fluorescence trace. Several recent studies have addressed this problem (Jewell & Witten 2018, Jewell et al. 2019, Friedrich et al. 2017, Pnevmatikakis et al. 2016). These studies have established a foundation for inferring spike times from fluorescence traces associated with a single neuron's activity (typically recorded in two-photon cameras; as originally phrased by (Vogelstein et al. 2009)). These studies utilized the dynamics of the calcium indicator concentration $\hat{c}$, its relation to the observed fluorescence $\hat{y}$, and its dependency on spike occurrences $s$ to phrase the following auto-regression model:

$$\begin{aligned} &\hat{y}_t^i = \hat{\beta}_0^i + \hat{\beta}_1 c_t^i + \hat{\epsilon}_t^i, \qquad \hat{\epsilon}_t^i \sim_{\text{ind.}} \left(0, (\hat{\sigma}^i)^2\right), \quad t = 1, \ldots, T, \\ &\hat{c}_t^i = \gamma \hat{c}_{t-1}^i + s_t^i, \quad t = 2, \ldots, T, \end{aligned} \tag{13}$$

In the model described above, the observed fluorescence of a neuron $i$ at timepoint $t$ , denoted as $\hat{y}_t^i$, is determined by the neuron's calcium concentration at that timepoint, $\hat{c}_t^i$ up to some factor $\hat{\beta}_1$, a constant shift, $\hat{\beta}_0$, and noise $\hat{\epsilon}_t^i$, which is assumed to be Gaussian and i.i.d.. In other words, the fluorescence is a noisy linear readout of the calcium concentration. The calcium concentration at each timepoint, $\hat{c}_t^i$, is the result of its decayed value from the previous timepoint, $\hat{c}_{t-1}^i$, by a factor $\gamma \in (0, 1)$ and the number of spikes emitted at that time $s_t^i$. In other words, the calcium concentration decays exponentially over time according to its decay constant and "jumps" with each spike emission.

We model the dynamics of a fluorescence trace arising from the activity of many neurons by summing the equations above across the population of neurons that contribute to the recorded fluorescence. This approach assumes that the camera operates within its linear dynamic range, so the recorded signal is a linear function of the incoming light. As a result, the total fluorescence is the linear sum of the individual neuronal fluorescence signals. For mathematical convenience, we divide this sum by the number of neurons contributing to it, $N$. Although $N$ is unknown, it is more practical to work with average quantities rather than total sums. In the resulting model, $N$ appears only implicitly, so its unknown value does not hinder our analysis. We assume that all neurons share the same calcium indicator dynamics, characterized by a similar decay constant $\gamma$, and similar relationships to fluorescence emission. We discuss this assumption in detail below. This leads us to the following autoregressive model for a fluorescence trace generated by the activity of many neurons, as appears in the main text, equation (1):

$$\begin{aligned} & y_t = \beta_0 + \beta_1 c_t + \epsilon_t, \qquad \epsilon_t \sim_{\text{ind.}} \left(0, \sigma^2\right), \quad t = 1, \ldots, T, \\ & c_t = \gamma c_{t-1} + r_t, \quad t = 2, \ldots, T. \end{aligned}$$

where $y_t \equiv \frac{1}{N}\sum_{i=1}^{N} \hat{y}_t^i$ is the total observed fluorescence from all $N$ neurons at timepoint $t$. $\beta_0 \equiv \frac{1}{N}\sum_{i=1}^{N} \hat{\beta}_0^i$, $c_t \equiv \frac{1}{N}\sum_{i=1}^{N} \hat{c}_t^i$, and $\epsilon_t \equiv \frac{1}{N}\sum_{i=1}^{N} \hat{\epsilon}_t^i$ with $\sigma^2 = \sum_{i=1}^{N} \left(\frac{\hat{\sigma}^i}{N}\right)^2$ are the total baseline fluorescence, total calcium at timepoint $t$, and total noise with total variance, respectively.

The population spiking rate, $r_t = \frac{1}{N}\sum_{i=1}^{N} s_t^i$, is the total number of spikes emitted at time $t$ by all $N$ neurons whose activity is contributing to the recorded fluorescence. Introducing the spiking rate into the model marks a significant change from the single-neuron model (13). With a single neuron, the number of spikes at each time point $s_t^i$ is a sparse quantity, with many zero entries and occasional small single-digit values. On the other hand, with many neurons, we expect the spiking rate to be a larger number that changes gradually over time. Since both the number of spikes in the single-neuron model, $s_t^i$, and the spiking rate in the multi-neuron model, $r_t$, are at the heart of the pursued solutions, their different properties significantly influence the approach we take to inference. We discuss this in Section 3 in the main text.

We note that it is possible that not all neurons contribute equally to the recorded fluorescence. In this case, each neuron $i$ has its own relative contribution, which we denote as $\alpha^i$, where $\sum_{i=1}^{N} \alpha^i = 1$. Consequently, the total fluorescence is given by $y_t \equiv \sum_{i=1}^{N} \alpha^i \hat{y}_t^i$. And we can calculate its dynamics by summing the dynamics of all individual neurons contributing to it, while weighting each neuron's dynamics (13) by its corresponding contribution $\alpha^i$. This leads to the multi-neuron model in equation (1), where the spiking rate is given by $r_t = \sum_{i=1}^{N} \alpha^i s_t^i$. In this case, all other quantities are also expressed in terms of each neuron's relative contribution. The total baseline fluorescence is given by $\beta_0 \equiv \sum_{i=1}^{N} \alpha^i \hat{\beta}_0^i$, the total calcium at timepoint $t$ is $c_t \equiv \sum_{i=1}^{N} \alpha^i \hat{c}_t^i$, and the total noise is $\epsilon_t \equiv \sum_{i=1}^{N} \alpha^i \hat{\epsilon}_t^i$ with its total variance $\sigma^2 = \sum_{i=1}^{N} \left(\alpha^i \hat{\sigma}^i\right)^2$. A key point relevant to this case is that the weighted rate remains a positive number that gradually changes over time, and its relation to the total fluorescence is still described by equation (1). We use these weights to calculate the population spiking rate of the biophysical Naomi dataset in Section 5.3 in the main text.

### B Proofs of Mathematical Results

#### Proof of Proposition 1

**Proposition 1.** *The pair* $(\tilde{r}, \tilde{\beta}0)$ *is a solution to problem (3) if and only if* $\tilde{\beta}_0 = \frac{1}{T}\mathbb{1}^\top(y - D^{-1}\tilde{r})$ *and* $\tilde{r}$ *is a solution to*

$$\text{minimize}_{r_1,\ldots,r_T} |\tilde{y} - Ar|^2 + \lambda \sum_{t=3}^{T}(r_t - r_{t-1})^2 \text{ subject to } r_t \geq 0, ; t = 2, \ldots, T, \tag{14}$$

*where* $A = \left(I - \frac{1}{T}\mathbb{1}\mathbb{1}^\top\right) D^{-1}$ *and* $\tilde{y} = \left(I - \frac{1}{T}\mathbb{1}\mathbb{1}^\top\right) y$*, with* $I$ *the* $T \times T$ *identity matrix and* $\mathbb{1}\mathbb{1}^\top$ *a* $T \times T$ *matrix of ones.*

**Lemma 1.** *Let* $A = PD^{-1}$*,* $\tilde{y} = Py$ *with* $P = I - \frac{1}{T}\mathbb{1}\mathbb{1}^\top$ *and* $\beta_0 = \frac{1}{T}\sum_{t=1}^{T}(\overline{y}_t - (D^{-1}r)_t)$*. Then,* $\|y - \mathbb{1}\beta_0 - D^{-1}r\|^2 = \|\tilde{y} - Ar\|^2$*.*

*Proof of Lemma 1.* Since $P^\top P = P$ and $P\mathbb{1} = 0$, it follows that

$$\|y - \mathbb{1}\beta_0 - D^{-1}r\|^2 = \left(y - 1_T\beta_0 - D^{-1}r\right)^\top \left(I - \frac{1}{T}\mathbb{1}\mathbb{1}^\top + \frac{1}{T}\mathbb{1}\mathbb{1}^\top\right)\left(y - \mathbb{1}\beta_0 - D^{-1}r\right)$$

$$= \left(y - \mathbb{1}\beta_0 - D^{-1}r\right)^\top P^\top P \left(y - \mathbb{1}\beta_0 - D^{-1}r\right) + \left(y - \mathbb{1}\beta_0 - D^{-1}r\right)^T \frac{1}{T}\mathbb{1}\mathbb{1}^\top \left(y - \mathbb{1}\beta_0 - D^{-1}r\right)$$

$$= \|\tilde{y} - Ar\|^2 + \frac{1}{T}\|\mathbb{1}^\top y - \mathbb{1}^\top\mathbb{1}\beta_0 - \mathbb{1}^\top D^{-1}r\|^2 = \|\tilde{y} - Ar\|^2.$$

The last equality follows from the fact that $\mathbb{1}^\top y - \mathbb{1}^\top\mathbb{1}\beta_0 - \mathbb{1}^\top D^{-1}r = 0$. □

*Proof of Proposition 2.* Taking the derivative of (5) with respect to $\beta_0$ and setting it equal to zero, we find that $\beta_0 = \frac{1}{T}\sum_{t=1}^{T}(y_t - (D^{-1}r)_t)$. The result follows from Lemma 1. □

### C Proof of Proposition 2

**Proposition 2.** *Let* $\bar{r}$ *denote a solution to optimization problem (7). Then, for any constant number* $d$*,* $\tilde{r} = \bar{r} + d\mathbb{1} + \tilde{r}_{m1}$ *also solves (7) where* $\tilde{r}_{m1}^\top = (r_{m1}, 0, 0, \ldots, 0)$ *is a* $T$*-length vector with all entries equal zero except for its first entry which satisfies*

$$r_{m1} = -d\frac{\sum_{j=1}^{T} A_{1j}}{A_{11}} \tag{15}$$

**Lemma 2.** *For any* $i = 1, \ldots, T$ *the following equality holds* $\frac{\sum_{j=1}^{T} A_{i,j}}{A_{i,1}} = \frac{\sum_{j=1}^{T} A_{1,j}}{A_{1,1}}$

*Proof of Lemma 2.* Equation(2) implies that

$$c_t = \sum_{t'=1}^{t} \gamma^{t-t'} r_{t'}. \tag{16}$$

Since $c = D^{-1}r$, we find from (16) that the entries of $D^{-1}$ are given by

$$\left(D^{-1}\right)_{i,j} = \begin{cases} 0, & \text{if } i < j \\ 1, & \text{if } i = j \\ \gamma^{i-j}, & \text{if } i > j \end{cases}. \tag{17}$$

From (17) we find the sum of the elements in each $D^{-1}$ row $i$,

$$\sum_{j=1}^{T} \left(D^{-1}\right)_{i,j} = \sum_{j=1}^{T} \gamma^{j-1} = \frac{1-\gamma^i}{1-\gamma}, \tag{18}$$

where the last step was calculated using the expression for the sum of a geometric series. Similarly for the columns,

$$\sum_{j=1}^{T} \left(D^{-1}\right)_{j,i} = \sum_{j=1}^{T-i+1} \gamma^{j-1} = \frac{1-\gamma^{T-i+1}}{1-\gamma}. \tag{19}$$

Recall that $A = \left(I - \frac{1}{T}\mathbb{1}\mathbb{1}^\top\right) D^{-1}$, with $I$ the identity matrix and $\mathbb{1}\mathbb{1}^\top$ a matrix of ones, is the column means subtracted version of $D^{-1}$. Hence,

$$\sum_{j=1}^{T} A_{i,j} = \sum_{j=1}^{T} \left(D^{-1}\right)_{i,j} - \sum_{j=1}^{T} \frac{1}{T} \sum_{k=1}^{i} \left(D^{-1}\right)_{k,j} = \frac{1-\gamma^i}{1-\gamma} - \sum_{j=1}^{T} \frac{1}{T} \frac{1-\gamma^{T-j+1}}{1-\gamma}, \tag{20}$$

where the last step was calculated using (18) and (19). We evoke once again the expression for the sum of a geometric series, and rearrange the result to receive from the above equation the following,

$$\sum_{j=1}^{T} A_{i,j} = -\frac{\gamma}{(1-\gamma)} \left(\gamma^{i-1} - \frac{1}{T}\frac{1-\gamma^T}{1-\gamma}\right). \tag{21}$$

The structure of $A$ and $D^{-1}$ column sums (19) also gives the following,

$$A_{i,1} = -\left(D^{-1}\right)_{i,1} - \frac{1}{T} \sum_{k=1}^{i} \left(D^{-1}\right)_{k,1} = \gamma^{i-1} - \frac{1}{T}\left(\frac{1-\gamma^T}{1-\gamma}\right). \tag{22}$$

From (22) and (21) we have,

$$\frac{\sum_{j=1}^{T} A_{i,j}}{A_{i,1}} = -\frac{\gamma}{(1-\gamma)}. \tag{23}$$

This implies that $\frac{\sum_{j=1}^{T} A_{i,j}}{A_{i,1}}$ is independent of $i$, including $i = 1$, and therefore one can write for every $i$, $\frac{\sum_{j=1}^{T} A_{i,j}}{A_{i,1}} = \frac{\sum_{j=1}^{T} A_{1,j}}{A_{1,1}}$

□

*Proof of Proposition 2.* Lemma 2 implies that $Ad\mathbb{1} + A\tilde{r}_{m1} = 0$ for any $d$ constant number and $\tilde{r}_{m1}$ a $T$-length vector with all entries equal zero except for its first entry which satisfies $r_{m1} = -d\frac{\sum_{j=1}^{T} A_{1j}}{A_{11}}$. Therefore,

$$\|\tilde{y} - A\tilde{r}\|^2 = \|\tilde{y} - (A\bar{r} + Ad\mathbb{1} + A\tilde{r}_{m1})\|^2 = \|\tilde{y} - A\bar{r}\|^2. \tag{24}$$

By construction of $\tilde{r}$ and $\tilde{r}_{m1}$, $\tilde{r}$ also satisfies

$$\sum_{t=3}^{T}(\tilde{r}_t - \tilde{r}_{t-1})^2 = \sum_{t=3}^{T}(\bar{r}_t - \bar{r}_{t-1})^2. \tag{25}$$

The convexity of problem (7) ensures that if $\bar{r}$ solves it, and $\tilde{r}$ maintains the value of its objective, as the previous two equations implies, than $\tilde{r}$ is also a solution to this problem.

□

### D Simulating spiking rate traces

We generated continuous spiking rate traces by integrating the following neural network $\frac{dx}{dt} = -x + gJ\tanh(x)$ where $x$ is a vector of size $N = 1000$, $J$ is an $N \times N$ matrix with i.i.d. entries $J_{i,j} \sim N(0, N)$, and we varied $g$ between 3.1 and 11.4 for the desired aimed fluctuations span $0.04 \ldots 0.35$ (Figure 2c; for Figure 2a&b we used $g = 5.7$ that generated 0.13 fluctuations level). We integrated with steps $dt = 0.05$ where each step represents 10ms of neural activity. These choices led to chaotic dynamics (Sompolinsky et al. 1988). To ensure non-negativity of the rate traces, we subtracted their minimum activity.

### E Clancy et al. Supplemental Figure 4 Dataset

Clancy et al. (2019) recorded wide-field fluorescence and, in parallel, spikes originating from the exact physical location as the fluorescence. The recording was done in V1. It includes a single fluorescence trace and a single spike counts trace, each with 130000 measurement points, taken at a 40hz rate (portraying almost an hour of activity). This dataset was obtained to verify that the fluorescence reflects the neural activity (to first order). Clancy et al. (2019) concluded it does so; see Clancy et al. (2019) Supplemental Figure 4c. They have reported to us a calcium decay of $\gamma = 0.973$ (it does not appear in their manuscript) by directly recording single neuron fluorescence traces and their spikes, from the same experiment preparation.

Since the calcium indicator rise time is about 40ms, which is longer than the original time bin of the recording (of 25ms), we worked with a two-point average of the data (making the rise time shorter than the 50ms after-average-bin). This also significantly improved the signal-to-noise ratio. Our analyzed signal hence has a 20hz rate. We fitted the decay rate to $\gamma^2$ for the algorithms accordingly. We further divided the single long fluorescence and spike count traces into 65 traces each; this way, we generated 65 data "trials," each with 1000 measurement points (50sec long), resembling a possible typical neuroscience experiment.

Recall that recording spike counts originating from the exact location of the fluorescence does not guarantee a true underlying spiking rate of the fluorescence, only an estimate. While Clancy et al. (2019) already verified that this estimate is reliable, Our results also support this conclusion. This can be seen visually in Figures 1 & 5(a) where examples of the recorded fluorescence, their inference, and the recorded spiking rate dynamics highly agree. It is also seen in Figure 5(d), where fluorescence fluctuations are reasonably correlated with the recorded spiking rate fluctuations, meaning the recorded spikes indeed drive the fluorescence recorded activity.

### F Estimating $\beta_1$ from the data

To evaluate $\beta_1$ we first allow it to take any arbitrary value in equation (1) (and do not set it to be equal to one). We repeat the steps of writing the model in equation (1) in its matrix form and subtracting its parameter means. We receive the following relation (up to noise contributions):

$$\tilde{y} = \beta_1 A r. \tag{26}$$

To find $\beta_1$ we minimize the squared error of the above relation w.r.t. $\beta_1$,

$$\underset{\beta_1}{\text{minimize}}\{\|\tilde{y} - \beta_1 A r\|^2\} \tag{27}$$

yielding

$$(Ar)^\top \tilde{y} = \beta_1 (Ar)^\top A r, \tag{28}$$

and hence the solution

$$\beta_1 = (Ar)^\top \tilde{y} \left((Ar)^\top A r\right)^+, \tag{29}$$

where $\left((Ar)^\top Ar\right)^+$ is the pseudo inverse of $(Ar)^\top Ar$.

To estimate $\beta_1$ from data, we calculate $\beta_1$ according to equation (29) using the recorded spiking rate $r$ and the recorded fluorescence $y$ (recorded in parallel), with $\tilde{y} = Py$, $Ar = PD^{-1}r$, $P = I - \frac{1}{T}\mathbb{1}\mathbb{1}^\top$ and $D^{-1}$ as in equation (17). If multiple segments of recorded data exist from the same experiment or setup (the more the merrier) we average across all $\beta_1$ calculated from the different segments, while excluding outliers, to give a single value $\beta_1$ estimate.

### G Choosing Penalty $\lambda$

Adopted from Jewell et al. (2019), Algorithm 3.

---

**Algorithm 2** A cross-validation scheme for choosing $\lambda$

---

Given the fluorescence signal $y$, the calcium decay $\gamma$, and a set of potential penalties $\lambda_1 \dots \lambda_M$, do:

1. Repeat twice:
   (a) Assign odd timesteps of the signal to the training set, $y^{train}$, and even timesteps to the test set, $y^{test}$, for the first repeat, and vice versa for the second repeat.
   (b) With each $m = 1 \dots M$ do:
      i. Solve $r_m^{train}$ with Con-Var (Algorithm 1) using $y^{train}$, $\gamma^2$ and $\lambda_m$.
      ii. Convolve $c_m^{train} = D^{-1} r_m^{train}$.
      iii. Average adjacent values in order to obtain predictions on the test set $\hat{c}_m^{test} = \frac{c_{m;1:T/2-1}^{train} + c_{m;2:T/2}^{train}}{2}$
      iv. Calculate the test set MSE for current repeat $i$, $ycMSE_m^i = \frac{2}{T} \sum_{t=1}^{T/2} (y^{test} - \hat{c}^{test})^2$
2. Average the test set MSE over repeats for each m yielding $\overline{ycMSE}_m$.
3. Find $\hat{m} = \text{argmin}_m \{\overline{ycMSE}_m\}$
4. Calculate the standard error of the test set MSE over repeats, for each m, $se(ycMSE)_m = \sqrt{\frac{\left(\overline{ycMSE}_m^1 - \overline{ycMSE}_m\right)^2 + \left(\overline{ycMSE}_m^2 - \overline{ycMSE}_m\right)^2}{2}}$
5. Find $m* = \max\{m : \overline{ycMSE}_m \leq \overline{ycMSE}_{\hat{m}} + se(ycMSE)_{\hat{m}}\}$
6. Return $\lambda_{m*}$

---
